# Cytotoxic lymphocytes use mechanosurveillance to target biophysical vulnerabilities in cancer

**DOI:** 10.1101/2020.04.21.054304

**Authors:** Maria Tello-Lafoz, Katja Srpan, Jing Hu, Yevgeniy Romin, Annalisa Calò, Katharine C. Hsu, Joan Massagué, Morgan Huse, Ekrem Emrah Er

## Abstract

Immune cells identify cancer cells by recognizing characteristic biochemical features indicative of oncogenic transformation. Cancer cells have characteristic mechanical features, as well, but whether these biophysical properties also contribute to destruction by the immune system is not known. In the present study, we found that enhanced expression of myocardin related transcription factors (MRTFs), which promote migration and metastatic invasion, paradoxically compromised lung colonization by melanoma and breast carcinoma cells in an immune-mediated manner. Cancer cells with increased MRTF signaling were also more sensitive to immune checkpoint blockade therapy in mice and humans. The basis for this vulnerability was not biochemical, but biophysical. MRTF expression strengthened the actin cytoskeleton, increasing the rigidity of cancer cells and thereby making them more vulnerable to cytotoxic T lymphocytes and natural killer cells. These results reveal a mechanical dimension of immunosurveillance, which we call mechanosurveillance, that is particularly relevant to the targeting of metastatic disease.

## INTRODUCTION

Immune cells detect and eliminate cancer cells by recognizing characteristic features that are indicative of oncogenic transformation. This process, known as immunosurveillance, is critical for the destruction of incipient neoplastic growth and plays a central role in anti-cancer immunotherapy (Finn, 2018). It is generally thought that immunosurveillance is mediated by molecular cues, such as stress-ligands, neoantigens, and danger associated molecular patterns, that trigger activating receptors on patrolling immune cells (Hernandez et al., 2016; Schumacher et al., 2019; Vesely et al., 2011). These biochemical features, however, are not exhibited by all cancer cells at all stages of disease, and they can be found in untransformed tissue, as well. Hence, effective immunosurveillance must utilize additional determinants. In that regard, it is intriguing that cancer progression also involves profound changes in cellular architecture and mechanics (Hall, 2009; Northcott et al., 2018; Suresh, 2007). These biophysical events are critical for promoting migration and invasive capacity, but whether they also serve as a basis for immunosurveillance is not known.

Cytotoxic lymphocytes, comprising natural killer (NK) cells and cytotoxic T lymphocytes (CTLs), play a central role in anti-cancer immunosurveillance by engaging and destroying transformed cells (Finn, 2018). Their cytolytic activity is initially triggered by recognition of surface molecules characteristic of stress and transformation, including cognate peptide-major histocompatibility complex (MHC) and the UL16 binding proteins, which engage the T cell antigen receptor (TCR) and the activating NK receptor NKG2D, respectively (Lanier, 2005; Zhang and Bevan, 2011). The binding of these and other stimulatory ligands drives the formation of a stereotyped interface between the lymphocyte and its target, called the immune synapse (Dustin and Long, 2010) The lymphocyte then secretes toxic granzyme proteases and the pore forming protein perforin into the synaptic space, thereby inducing target cell apoptosis.

Immune synapses are physically active structures, exerting nanonewton scale mechanical forces that enhance the efficiency of perforin and granzyme-mediated killing (Bashour et al., 2014; Basu et al., 2016; Husson et al., 2011). These forces are also thought to facilitate lymphocyte activation by promoting mechanotransduction. Indeed, several activating immunoreceptors, including the TCR, only reach full signaling capacity under applied force (Friedland et al., 2009; Liu et al., 2014). This requirement places physical demands on the target cell surface, which must presumably be rigid enough to counterbalance the mechanical load placed upon receptor-bound ligands. Consistent with this idea, stiff surfaces bearing stimulatory ligands induce substantially stronger lymphocyte activation than softer surfaces coated with the same proteins (Blumenthal et al., 2019; Comrie et al., 2015; Judokusumo et al., 2012; Saitakis et al., 2017; Wan et al., 2013). Hence, it is not unreasonable to expect that the biophysical properties of cancer cells might regulate their susceptibility to cytotoxic lymphocyte-mediated attack.

Cytotoxic lymphocytes are particularly effective at combatting metastatic cancer cells (Dye, 1986; Eyles et al., 2010; Malladi et al., 2016; Pommier et al., 2018; Wei et al., 2018), which live alone or in small groups far from the immunosuppressive microenvironment of the primary tumor. Interestingly, metastasis is associated with dramatic morphological and biophysical change, mostly driven by remodeling of the filamentous actin (F-actin) cytoskeleton. This supports multiple steps in the metastatic cascade, including local invasion from the primary tumor, intravasation into circulation, and subsequent extravasation into target organs (Bravo-Cordero et al., 2012). It is now becoming clear that F-actin dynamics are also critical for metastatic outgrowth in the new microenvironment, which typically occurs in the perivascular niche, a nutrient rich milieu on the abluminal surface of microvessels (Ghajar et al., 2013; Kienast et al., 2010). To expand successfully in this space, metastatic cells first establish strong adhesion to the microvascular basement membrane (Shibue and Weinberg, 2009; Valiente et al., 2014). This triggers a mechanotransduction response in which cell spreading and migration are coupled to the activation of myocardin-related transcription factors (MRTF) A and B (Er et al., 2018). In the steady state, MRTF isoforms are sequestered in the cytoplasm via binding to monomeric globular actin (G-actin). Cytoskeletal growth, often induced by Rho-family GTPases like RAC1, depletes this G-actin pool, liberating MRTFA and MRTFB to enter the nucleus (Gau and Roy, 2018; Gualdrini et al., 2016; Kim et al., 2017; Lionarons et al., 2019; Medjkane et al., 2009; Olson and Nordheim, 2010). Once localized in this manner, MRTFs form complexes with the DNA-binding protein serum response factor (SRF) to drive expression of more G-actin and cytoskeletal components. Morphoregulation by MRTF is absolutely required for metastatic invasion and subsequent proliferative expansion (Er et al., 2018). However, because analogous shape changes have been shown to increase cell stifffness (Kasza et al., 2009), it is tempting to speculate that MRTF signaling might also mechanically sensitize cancer cells to cytotoxic lymphocytes.

In the present study, we explored this hypothesis by analyzing the effects of MRTF on the mechanical properties of cancer cells, their immune sensitivity, and their capacity to colonize tissues *in vivo*. Our results reveal a novel, mechanical form of immunosurveillance that enables cytotoxic lymphocytes to target the specific biophysical features of metastatic cancer cells.

## RESULTS

### MRTF overexpression sensitizes metastatic cells to the immune system

Given the importance of MRTF-mediated mechanotransduction for migration and metastasis, and the unique position of MRTF in F-actin regulation, we reasoned that manipulating MRTFA and MRTFB levels might reveal novel vulnerabilities associated with cancer cell architecture (Fig. 1A). To explore this idea, we employed an established model of metastatic colonization in which malignant cancer cells expressing luciferase are injected intravenously into congenic mice and their subsequent growth in the lung monitored by bioluminescent imaging (Fig. 1B). shRNA-mediated suppression of MRTFA and MRTFB reduced lung colonization by both B16F10 melanoma and E0771 breast cancer cells in this model (Fig. 1C-D and Fig. S1A), consistent with previous data showing that MRTF is required for metastatic seeding (Er et al., 2018; Kim et al., 2017; Medjkane et al., 2009). However, B16F10 and E0771 cell lines overexpressing MRTFB (B16F10-MRTFB and E0771-MRTFB) exhibited dramatically reduced lung colonization (Fig. 1E-F and Fig. S1A), the opposite of what one would expect for cells with enhanced MRTF-induced invasiveness. E0771-MRTFA cells also metastasized poorly, although B16F10-MRTFA cells displayed increased colonization (Fig. S2A).

**Fig. 1.**
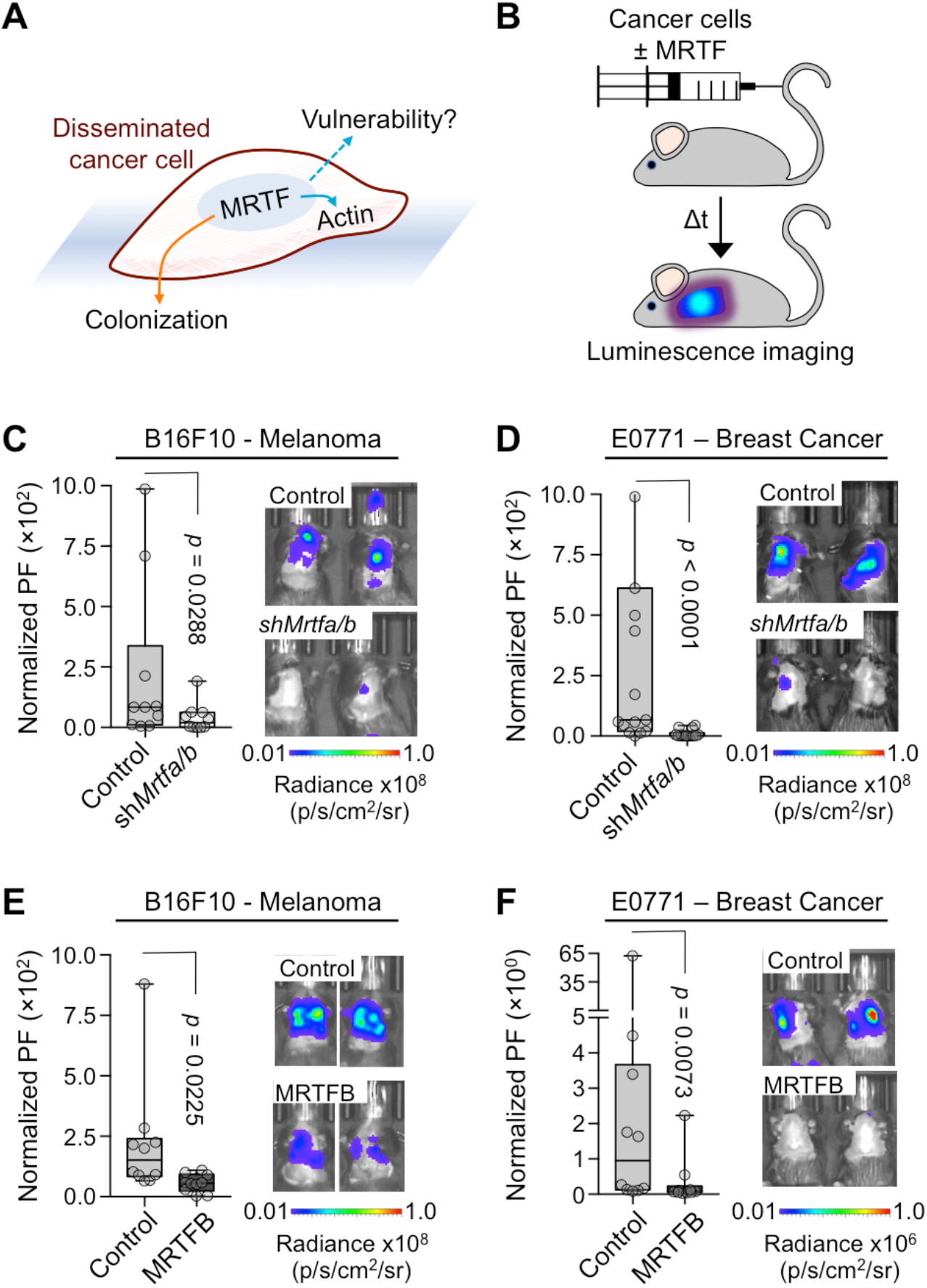
MRTF overexpression inhibits lung metastasis. (A) MRTF signaling promotes metastatic colonization but may also create vulnerabilities. (B) Experimental design for lung colonization model. (C-D) Metastatic burden in lungs of C57BL/6J mice injected with syngeneic control or *Mrtfa/b* knockdown B16F10 (C) or E0771 (D) cells, measured by bioluminescent imaging (BLI) 3 weeks after tail vein injection and normalized to the first day of injection. PF: photon flux (*n* = 10 mice per group). (E-F) BLI of mice 3 weeks post tail vein injection with B16F10 (E) or E0771 (F) cells overexpressing MRTFB or empty vector control (*n* = 10 mice per group). Box plots show upper and lower quartiles, median, maximum, and minimum values. *p* values were calculated by Mann-Whitney test. See also Fig. S1 and S2.

We initially hypothesized that elevated MRTF signaling might impair metastasis by inhibiting cancer cell proliferation. Overexpression of MRTFA and MRTFB, however, did not significantly alter cancer cell growth and division *in vitro*, as assessed by the acquisition of vital dye and the dilution of tracing stain, respectively (Fig. S1B-C). Proliferation *in vivo*, which we quantified by staining tumor sections for the Ki67 marker, was similarly unchanged (Fig. S1D). We also examined the effects of MRTF overexpression on steady state cell death, and found that only E0771-MRTFA cells exhibited increased apoptosis in isolation (Fig. S1E). These results suggested that the reduced metastatic capacity conferred by MRTF was not entirely intrinsic to the tumor cells and instead required component(s) of the metastatic microenvironment.

Histological analyses of B16F10 and E0771 lung lesions revealed a substantial number of infiltrating CD8^+^ CTLs and NK cells (Fig. 2A and Fig. S2B). To assess the importance of these lymphocytes for MRTF dependent cancer cell death, we depleted mice of NK cells and CTLs using anti-asialo GM1 and anti-CD8 antibodies, respectively, prior to injecting B16F10 or E0771 cells (Fig. 2B). NK cell depletion dramatically enhanced colonization by B16F10-MRTFA and B16F10-MRTFB cells (Fig. 2C and Fig. S2C). In the case of B16F10-MRTFB cells, this effect led to a striking phenotypic reversal in which the overexpressing cells now exhibited stronger metastatic growth than B16F10 controls (compare Fig. 1E to 2C). Depletion of CD8^+^ T cells induced similarly remarkable increases in both E0771-MRTFB and E0771-MRTFA colonization (Fig. 2D and Fig. S2D). Interestingly, NK depletion did not rescue the metastatic activity of E0771-MRTFB cells (Fig. S2E), implying that these cells were primarily constrained by CTLs *in vivo*, whereas B16F10-MRTFB tumors were subject to NK-mediated control. Collectively, these results demonstrate that the increased metastatic potential conferred by MRTF expression is curbed by cytotoxic lymphocytes (Fig. 2E).

**Fig. 2.**
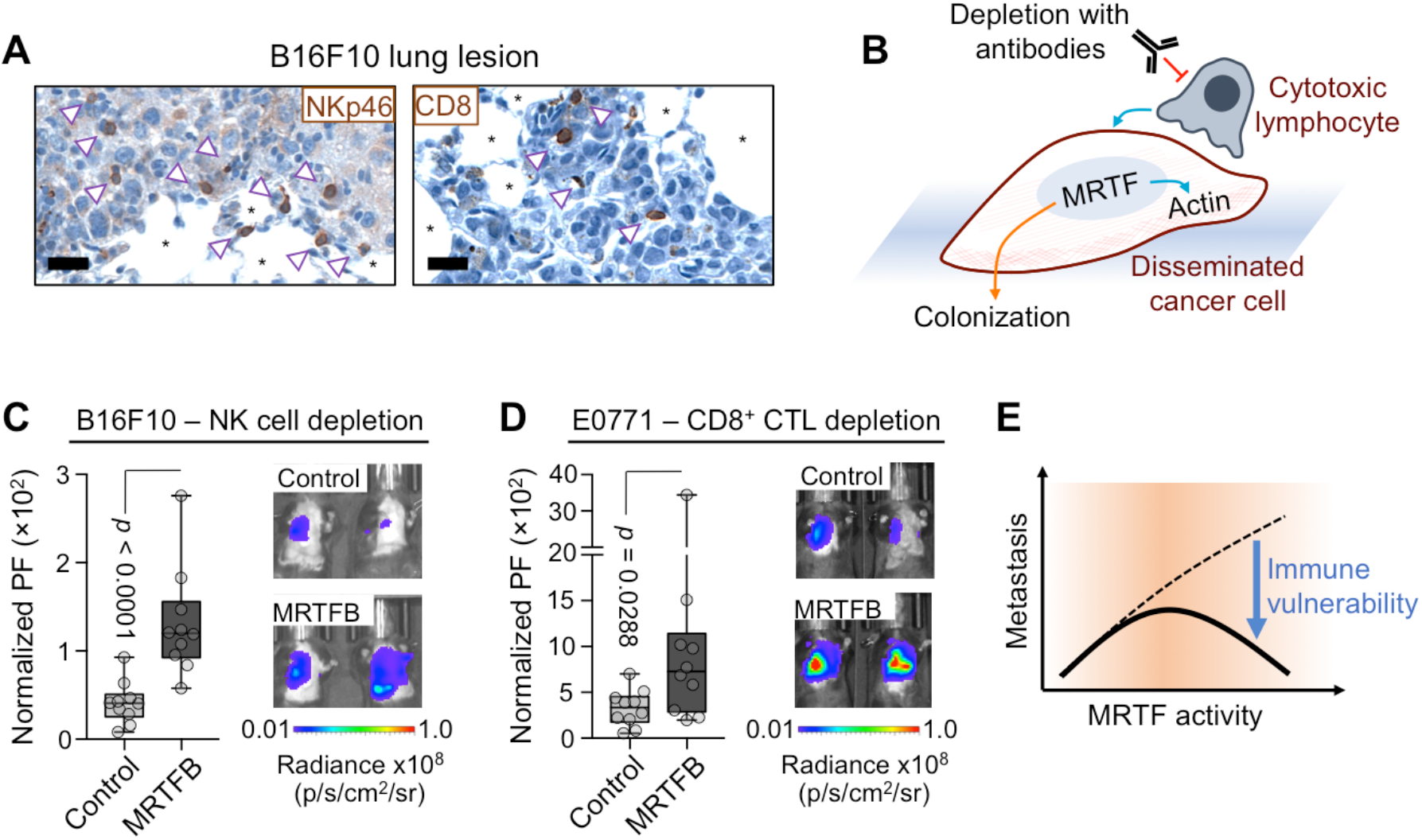
MRTF sensitizes cells to cytotoxic lymphocytes. (A) Representative IHC images of NK cell (arrowheads, NKp46 staining) and CD8^+^ T cell (arrowheads, CD8 staining) infiltration in B16F10 lung metastases. *: alveolar space, scale bars: 20 μm. (B) Using blocking antibodies to inhibit immunosurveillance by cytotoxic lymphocytes. (C-D) BLI of mice pretreated with anti-asialo GM1 antibody (C) or anti-CD8 antibody (D) for NK and CD8^+^ T cell depletion, respectively, and imaged 2 weeks after injection of the indicated cancer cells (*n* = 10 mice per group). (E) Model showing correlation between increased MRTF activity and metastatic potential (dashed line). These properties are uncoupled because high MRTF expression sensitizes metastatic cells to cytotoxic lymphocytes (solid line). Box plots show upper and lower quartiles, median, maximum, and minimum values. *p* values were calculated by Mann-Whitney test. See also Fig. S1 and S2.

### MRTF renders cancer cells more stimulatory to cytotoxic lymphocytes

The importance of CTLs and NK cells for the MRTF dependent suppression of metastasis *in vivo* suggested that MRTF signaling might make cancer cells more vulnerable to cellular cytotoxicity. To investigate this hypothesis, we loaded B16F10 and E0771 cells containing different levels of MRTF with ovalbumin_257-264_ peptide (OVA) and then mixed them with CTLs expressing the OT1 TCR, which recognizes OVA bound to the MHC protein H2Kb. shRNA-induced suppression of MRTFA and MRTFB reduced B16F10 killing in these experiments, while overexpression of either MRTF isoform enhanced CTL-mediated lysis (Fig. 3A-B). E0771-MRTFA and E0771-MRTFB cells were also more sensitive to CTLs than E0771 controls (Fig. 3C). This effect was specific for cellular cytotoxicity, as overexpression of MRTFA and MRTFB did not, in general, boost apoptotic responses to staurosporine and tumor necrosis factor (TNF), although E0771-MRTFA cells were more vulnerable to these agents (Fig. S3A-B).

**Fig. 3.**
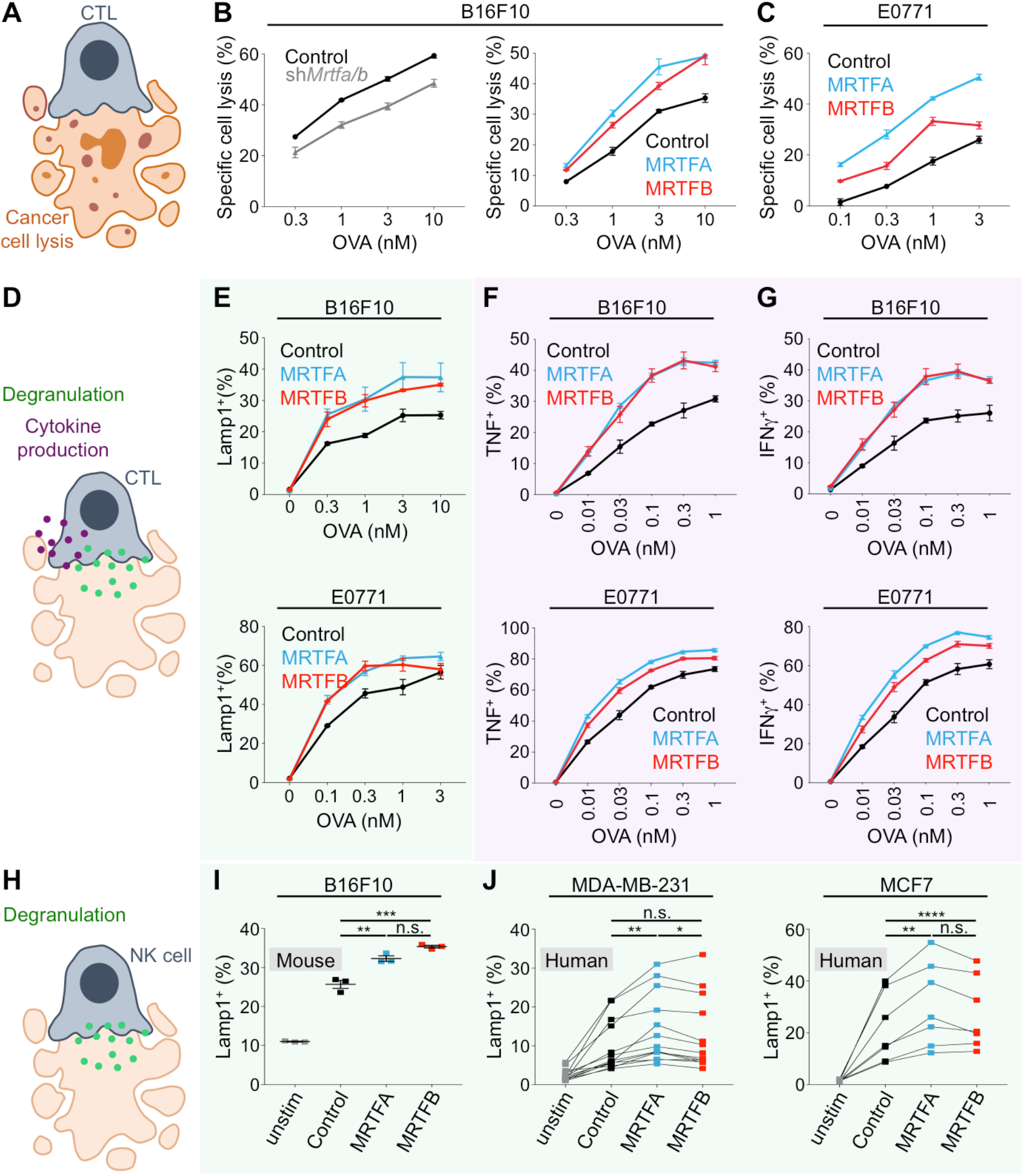
MRTF expression sensitizes cancer cells to CTL-mediated lysis. (A, D, H) Diagrams of CTL-mediated cancer cell lysis (A), CTL degranulation and cytokine secretion after cancer cell recognition (D), and NK cell degranulation after cancer cell recognition (H). (B-C, E-G) B16F10 or E0771 cell lines were loaded with increasing concentrations of OVA and mixed with OT1 CTLs. (B) Specific lysis of control or *Mrtfa/b* knockdown B16F10 cells (left) or B16F10 cells overexpressing MRTFA, MRTFB, or empty vector (right) 5 h after mixing with CTLs. (C) Specific lysis of E0771 cells overexpressing MRTFA, MRTFB, or empty vector (right) 5 h after mixing with CTLs. (E) Degranulation, measured by surface exposure of CTL Lamp1 90 min after mixing with B16F10 and E0771 cells overexpressing MRTFA, MRTFB, or empty vector. (F-G) Production of TNF (F) and IFN*γ* (G) measured by intracellular staining of CTLs 4 h after mixing with B16F10 and E0771 cells overexpressing MRTFA, MRTFB, or empty vector. (I) Splenic murine NK cells were mixed with B16F10 cells overexpressing MRTFA, MRTFB, or empty vector and degranulation quantified after 4 h. Data in B-C, E-G, and I are shown as mean ± SEM of technical triplicates, representative of 3 independent experiments. (J) Human NK cell clones derived from peripheral blood were mixed with the indicated control and MRTFA/B overexpressing cell lines. After 5 h, NK cell degranulation was measured by surface exposure of Lamp1. Black lines indicate samples derived from the same donor. (*n* = 9 donors for MCF7 experiments, *n* = 10 donors for MDA-MB-231). \*\*\*\**p* ≤ 0.0001, \*\*\**p* ≤ 0.001, \*\**p* ≤ 0.01, and \**p* ≤ 0.05, and n.s.: not significant for *p* > 0.05, calculated by one-way ANOVA. See also Fig. S3.

To investigate how MRTF signaling sensitizes cancer cells to cellular cytotoxicity, we exposed B16F10 and E0771 cell lines to purified perforin and granzyme B. Neither MRTF isoform increased cell death in these experiments (Fig. S3C), indicating that the intrinsic cellular response to perforin and granzyme was unchanged. Cytotoxic lymphocytes also produce Fas ligand (FasL), which induces apoptosis in target cells expressing the death receptor Fas (Nagata, 1999). Although E0771-MRTFA cells expressed elevated levels of Fas and underwent apoptosis in response to soluble FasL, E0771-MRTFB, B16F10-MRTFA, and B16F10-MRTFB cells expressed little to no Fas and were resistant to FasL-mediated killing (Fig. S3D-E). Hence, differential sensitivity to perforin/granzyme or FasL did not broadly explain how MRTF signaling makes cancer cells more vulnerable to cytotoxic attack.

Next, we examined whether cancer cells overexpressing MRTFA and MRTFB induce stronger lymphocyte activation. Perforin and granzymes are stored in specialized secretory lysosomes called lytic granules, which fuse with the plasma membrane after synapse formation (Stinchcombe and Griffiths, 2007). To quantify granule exocytosis, which is also called degranulation, we monitored surface exposure of the lysosomal marker Lamp1 in OT1 CTLs cocultured with antigen-loaded cancer cells (Fig. 3D). Antigen-loaded B16F10-MRTFA/B and E0771-MRTFA/B cells induced stronger degranulation responses than did their respective controls (Fig. 3E), implying that increased MRTF signaling in the target cell enables more effective CTL stimulation. Activated CTLs also generate and release the inflammatory cytokines interferon-γ (IFNγ) and TNF. Production of both of these cytokines was markedly enhanced in cocultures with MRTF overexpressing cancer cells (Fig. 3D, F-G). Collectively, these data indicated that MRTFA and MRTFB render cancer cells more stimulatory to cytotoxic lymphocytes.

To explore the generality of this paradigm, we extended our studies to splenic NK cells derived from C57BL/6 mice. Similar to our results with CTLs, we found that B16F10-MRTFA and B16F10-MRTFB cells induced significantly stronger NK cell degranulation than B16F10 controls (Fig. 3H-I). We also examined primary human NK cells, which recognize and destroy the breast carcinoma cells lines MDA-MB-231 and MCF7. Peripheral blood mononuclear cells (PBMCs, ~10% CD56^+^CD3^−^ NK cells) from multiple donors were mixed with parent MDA-MB-231 and MCF7 cells as well as lines overexpressing MRTFA and MRTFB. Degranulation responses were significantly stronger in cocultures with MRTF overexpressing cells (Fig. 3H, J), further supporting the interpretation that MRTF signaling boosts the stimulatory capacity of target cells. We conclude that MRTFA and MRTFB influence cytotoxic immune cell-cell interactions across both species and lymphocyte cell type.

### MRTF signaling enhances responsiveness to checkpoint blockade therapy

The capacity of MRTF signaling to enhance CTL responses against cancer cells implied that it might increase the efficacy of immune checkpoint blockade (ICB), a group of antibody-based immunotherapies that function by derepressing tumor specific T cells (Lesokhin et al., 2015; Wei et al., 2018). To investigate this possibility, we injected mice with control B16F10 or B16F10-MRTFB cells and then administered three doses of blocking antibody against CTLA4, a well-established inhibitory checkpoint receptor (Fig. 4A). Although anti-CTLA4 treatment moderately reduced lung colonization by control B16F10 cells, this effect failed to confer a survival benefit (Fig. 4B-C). By contrast, blocking CTLA4 significantly inhibited B16F10-MRTFB metastasis, leading to a marked increase in survival. These results indicate that MRTF signaling sensitizes cancer cells *in vivo* to a therapeutically enhanced immune system.

**Fig. 4.**
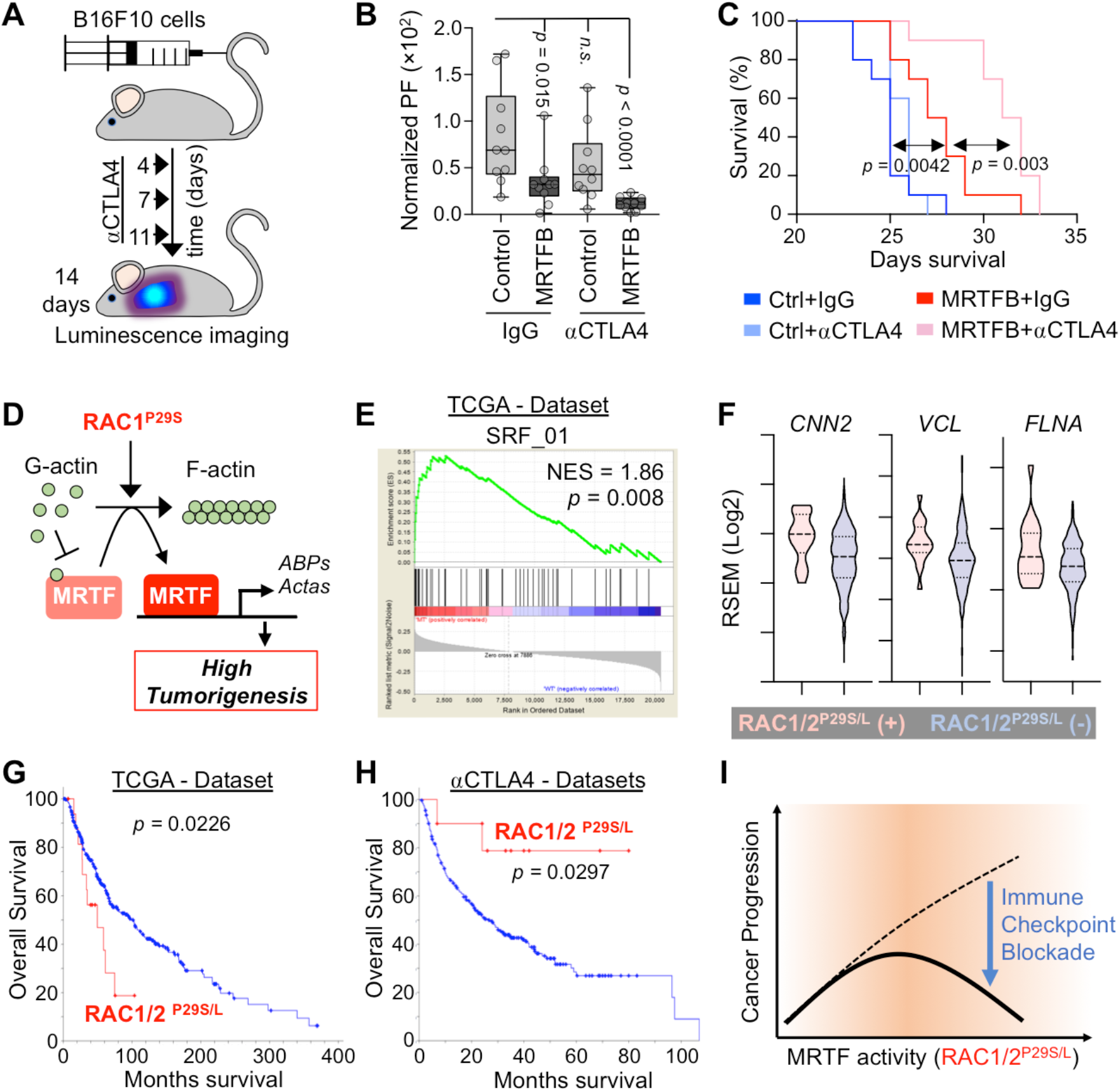
MRTF boosts therapeutic T cell responses in the context of anti-CTLA4 ICB. (A) Experimental design for anti-CTLA4 (αCTLA4) treatment of mice inoculated with B16F10 melanoma. (B) BLI of mice 2 weeks after injection with B16F10 cells overexpressing MRTFB or control vector and treatment with control IgG or anti-CTLA4 antibody (*n* = 10 mice per group). Box plots show upper and lower quartiles, median, maximum, and minimum values. *p* values were calculated using Mann-Whitney test, n.s.: not significant for *p* = 0.1655. (C) Kaplan-Meier survival curves for percent survival of mice in B (*n* = 10 mice per group). (D) Model showing oncogenic RAC1^P29S^ driven activation of MRTF, leading to increased tumorigenicity (Lionarons et al., 2019). *Acta:* actin family of proteins, *ABPs:* actin binding proteins. (E) GSEA showing MRTF-SRF target gene expression enrichment in RAC1/2^P29S/L^ mutant skin cutaneous melanoma patients in The Cancer Genome Atlas (TCGA) dataset. NES: normalized enrichment score. (F) Violin plots showing increased expression of known MRTF-SRF target genes in RAC1/2^P29S/L^ patients. Dashed lines medians, dotted lines upper and lower quartiles. Pink and blue plots represent data from patients with or without RAC1/2^P29S/L^ mutation, respectively. (G-H) Overall survival of RAC1/2^P29S/L^ patients in the TCGA dataset (G) and in melanoma patients treated with anti-CTLA4 ICB in a pooled dataset (H), which was derived from Samstein et al., 2019 (75 patients); Miao et al., 2018 (144 patients); Van Allen et al., 2015 (20 patients); Catalanotti et al., 2017 (21 patients), and Liang et al., 2017 (14 patients). (I) MRTF activity induced by RAC1/2^P29S/L^ potentiates melanoma progression (similar to Fig. 2E) but simultaneously sensitizes cancer cells to immune checkpoint blockade. *p* values in C, G, and H were calculated by Log-rank test. See also Fig. S4 and supplementary tables 1 and 2.

To further explore this hypothesis in the context of human ICB trials, we examined clinical data for links between MRTF signaling and responsiveness to anti-CTLA4 therapy. While there are no reported MRTF gain-of-function mutations in human solid tumors, MRTF signaling is strongly induced by the constitutively active P29S mutant form of the small GTPase RAC1 (Fig. 4D) (Lionarons et al., 2019). RAC1^P29S^ and related mutations (e.g. RAC1^P29L^) are found in ~5 % of human melanomas, a particularly aggressive subset that exhibits resistance to BRAF inhibitors in the clinic (Lionarons et al., 2019; Van Allen et al., 2014; Watson et al., 2014). Using the TCGA database, we were able to corroborate a link between these mutations and the MRTF-SRF pathway in human melanoma. Gene Set Enrichment Analysis (GSEA) revealed that RAC1/2^P29S/L^ tumors significantly upregulated genes containing SRF binding sites (Fig. 4E and Fig. S4A-B), including *CNN2* (calponin 2), *VCL* (vinculin), and *FLNA* (filamin A) (Fig. 4F). Based on our *in vivo* mouse experiments, we reasoned that these tumors might also be more sensitive to ICB. Analysis of TCGA data revealed that patients with RAC1/2^P29S/L^ melanoma exhibited reduced overall survival (Fig. 4G and Fig. S4B, Supplementary Table 1). In patients receiving anti-CTLA4 therapy, however, RAC1/2^P29S/L^ mutations correlated with significantly improved outcomes (Fig. 4H and Fig. S4D, Supplementary Table 2). Taken together, these results are consistent with the idea that MRTF signaling becomes a liability for tumor cells during anti-CTLA4 ICB by augmenting therapeutic T cell responses (Fig. 4I).

The RAC1^P29S^ mutation has been associated with UV damage (Cancer Genome Atlas, 2015), raising the possibility that the pro-survival effect we observed in the context of anti-CTLA4 (Fig. 4H) did not result specifically from MRTF signaling but rather from T cell recognition of UV-induced neoantigens. To explore this alternative hypothesis, we examined whether melanoma patients with UV-damage associated driver mutations other than RAC1/2^P29S/L^ also responded better to anti-CTLA4 therapy. UV-induced mutations in *PPP6C, IDH1,* and *FBXW7* did not correlate with increased survival, and while patients with *NF1* truncation did exhibit modestly improved responses, this effect was not statistically significant and, furthermore, it was primarily attributable to patients with co-occurring RAC1^P29S/L^ mutations (Fig. S4C-E). We also investigated the expression of MRTF-SRF signature genes, and found that melanomas with UV-induced *NF1, PPP6C, IDH1,* and *FBXW7* mutations failed to upregulate this gene set (Fig. S4D). We conclude that MRTF-SRF-induced gene expression, rather than UV damage alone, promotes increased responsiveness to anti-CTLA4 in RAC1/2^P29S/L^ melanoma.

### MRTF sensitizes cancer cells to cytotoxic lymphocytes by increasing stiffness

To investigate how MRTFA and MRTFB render cancer cells more stimulatory to cytotoxic lymphocytes, we performed whole transcriptome RNA-sequencing of B16F10 and E0771 cells overexpressing each transcription factor. MRTFA induced hundreds of expression changes in both cells lines, the majority of which were gene upregulation events (Fig. 5A-C). Some of the most strongly activated genes were actin isoforms and cytoskeletal regulators, among them *Acta1*, *Actg2*, *Fhl1*, and *Myh11*. Although MRTFB generated more modest expression changes, nevertheless it induced many of the same cytoskeletal genes (Fig. 5A-C), implying that it was these genes that were responsible for the shared effects of MRTFA and MRTFB on immune sensitization. To further explore this idea, we performed Gene Ontology analysis using gene sets induced by one MRTF isoform in both cells lines and also gene sets induced by both MRTF isoforms in each cell line (Fig. 5B Fig. S5). The results did not reveal substantial MRTF-induced expression of immune related pathways. Processes and components pertaining to the actin cytoskeleton and cellular architecture were dramatically induced, however, in line with the known functions of MRTF signaling (Gau and Roy, 2018; Olson and Nordheim, 2010) (Fig. 5B and Fig. S5). Consistent with these results, B16F10 and E0771 cells overexpressing MRTFA and MRTFB contained copious amounts of filamentous actin (F-actin), which in some cases formed dense arrays of stress fibers (Fig. 5D).

**Fig. 5.**
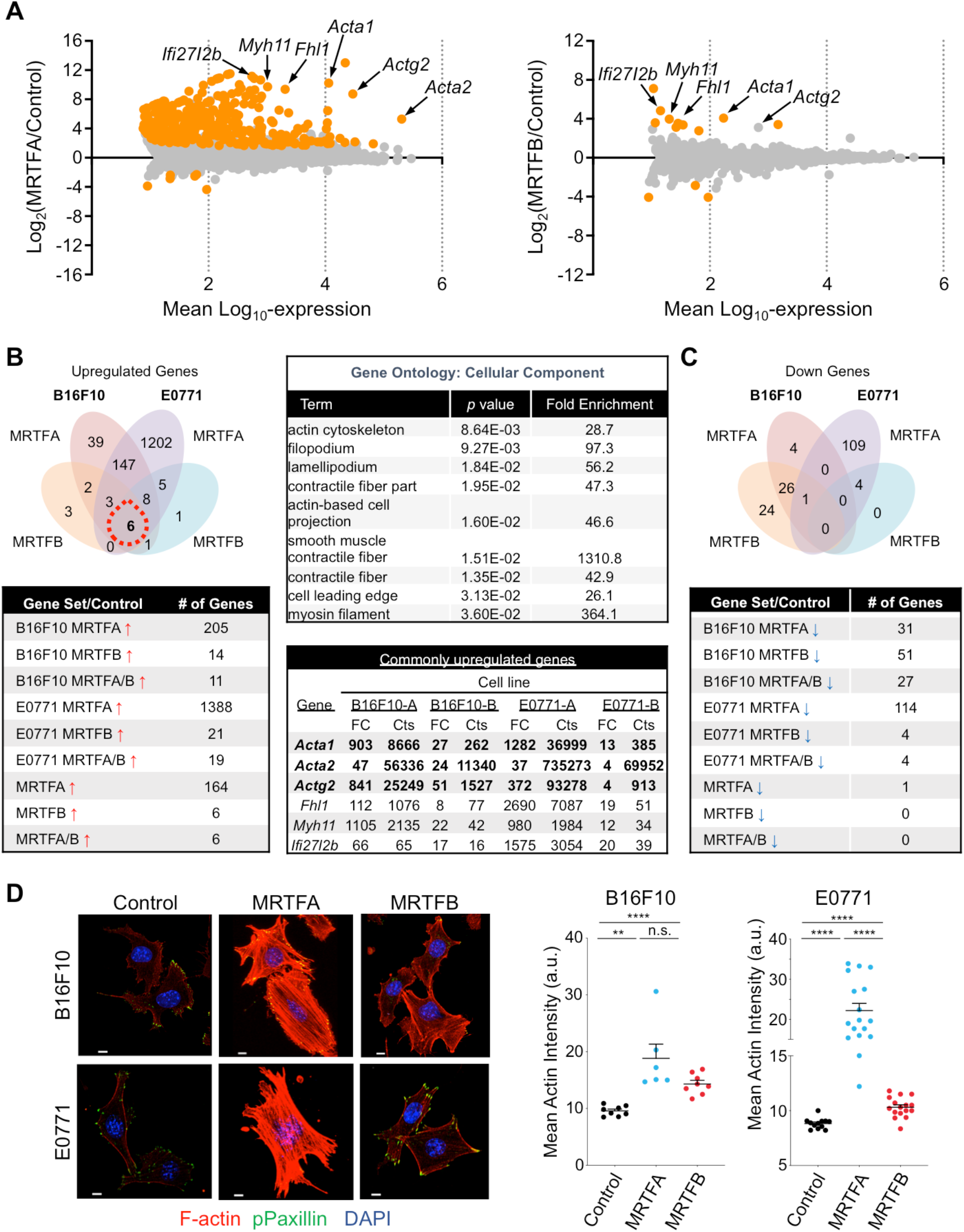
MRTF overexpression augments the F-actin cytoskeleton. (A) Mean-difference plots showing gene expression changes induced by MRTFA (left) and MRTFB (right). Graphs incorporate data from both B16F10 and E0771 cells. Statistically significant gene expression changes are colored orange (*p* ≤ 0.05, adjusted for multiple testing using the Benjamini method). Strongly upregulated genes encoding actin isoforms and F-actin regulators are indicated. (B) Above left, Venn diagram of upregulated genes exclusive to or shared by B16F10 and E0771 cells overexpressing MRTFA or MRTFB in culture. Below left, table showing the number of upregulated genes for each group. Above right, Gene Ontology (GO) analysis using the set of genes upregulated in all cell lines (red dashed circle in Venn diagram). Statistically significant GO terms are shown, with reported *p* values corrected for multiple testing using the Benjamini method. Below right, table of commonly upregulated genes, in which genes with over 50 RNA sequence counts in all data sets are shown in bold. FC: Fold Change, Cts: RNA sequencing read counts. (C) Above, Venn diagram of downregulated genes exclusive to or shared between B16F10 and E0771 cells overexpressing MRTFA or MRTFB in culture. Below, table showing the number of downregulated genes for each group. (D) Left, confocal images of representative B16F10 and E0771 control, MRTFA, and MRTFB overexpressing cells, stained with DAPI (blue), phalloidin (F-actin, red) and anti-phospho-paxillin (green). Scale bars: 10 μm. Right, graphs showing mean actin intensity for each cell line. Error bars denote SEM, n.s.: not significant for *p* > 0.05, \*\**p* ≤ 0.01, \*\*\*\**p* ≤ 0.0001; two-tailed paired Student’s t test; *n* ≥ 8 images per cell line; representative of 3 independent experiments. See also Fig. S5.

These striking architectural phenotypes raised the possibility that MRTF signaling might modulate immune activation by altering the biophysical properties of cancer cells. Given that the stiffness of the opposing surface controls activating mechanotransduction through the immune synapse (Blumenthal et al., 2019; Comrie et al., 2015; Judokusumo et al., 2012; Saitakis et al., 2017; Wan et al., 2013), and knowing the importance of the actin cytoskeleton for cellular architecture, we hypothesized that MRTF signaling might render cancer cells more stimulatory by increasing their rigidity (Fig. 6A). To investigate this hypothesis, we used atomic force microscopy (AFM)-based indentation to profile the effects of MRTFA and MRTFB on the deformability of B16F10, E0771, MDA-MB-231, and MCF7 cells (Fig. 6B). Overexpression of either MRTFA or MRTFB significantly increased the average and peak stiffness of every cell line examined (Fig. 6C-F). Stiffness measurements decreased dramatically in the presence of the F-actin depolymerizing agent latrunculin A (Fig. 6G), confirming the importance of the actin cytoskeleton for controlling this parameter (Rotsch and Radmacher, 2000). These results establish strong correlations between the biophysical effects of MRTF overexpression and the capacity of cancer cells to activate cytotoxic lymphocytes.

**Fig. 6.**
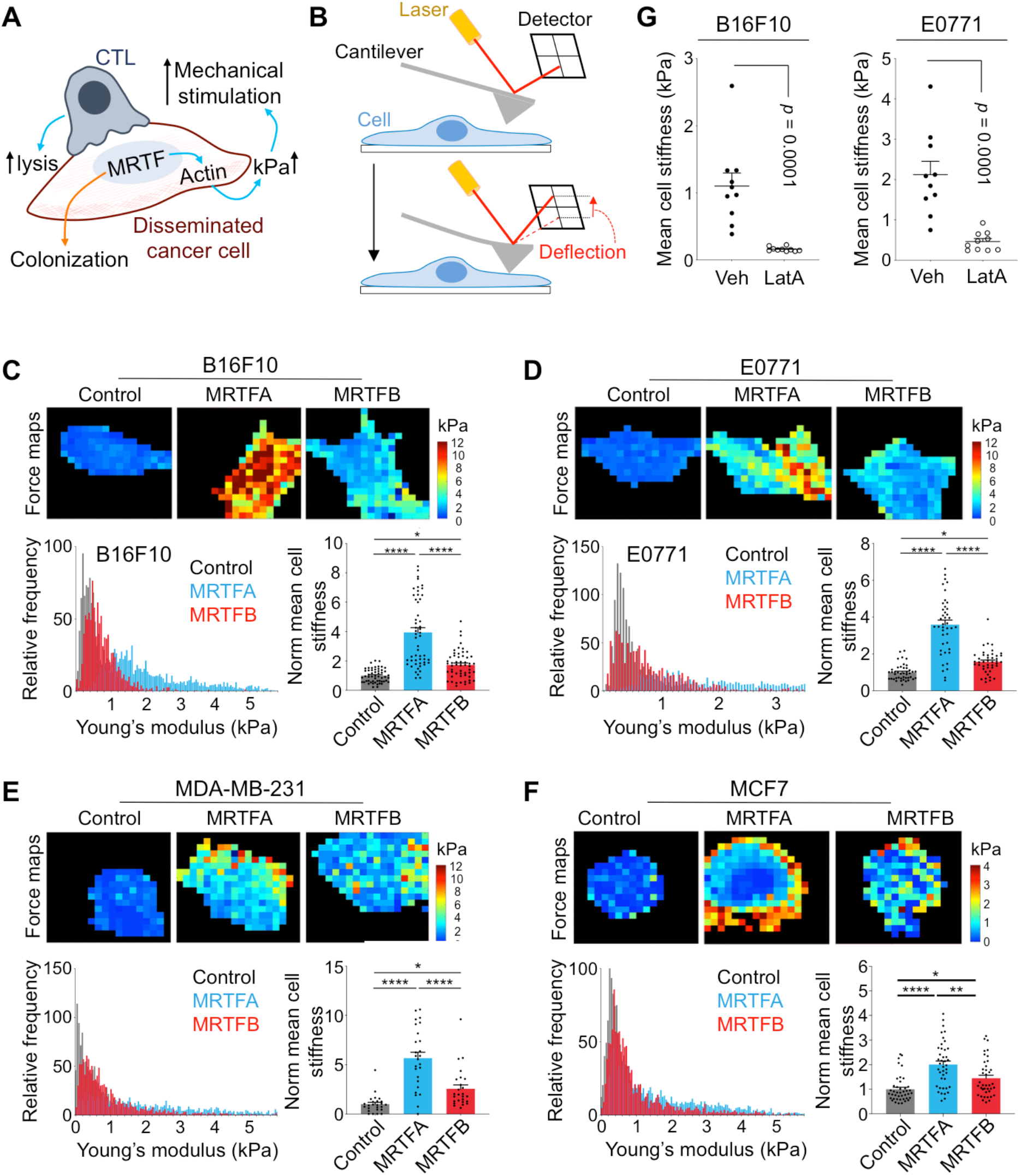
MRTF signaling increases cell stiffness. (A) MRTF activity promotes metastatic colonization but also sensitizes cancer cells to cytotoxic lymphocytes by boosting actin polymerization and thereby increasing cell stiffness (kPa). (B) Schematic diagram showing the AFM probe indentation approach. Cantilever deflection is proportional to the loading force and is used to calculate the Young’s modulus of a cell. (C-F) AFM stiffness measurements comparing B16F10 (C), E0771 (D), MDA-MB-231 (E), and MCF7 (F) cells overexpressing MRTFA or MRTFB with respective control cell lines. Above, force maps of representative cells, with Young’s modulus value (kPa) indicated in pseudocolor. Below left, probability histograms of pooled Young’s modulus measurements (kPa) from representative experiments, *n* = 10 cells. Below right, graphs of mean cell stiffness values normalized to the control. Data from 4 independent experiments are shown as mean ± SEM (One-way ANOVA with Tukey’s multiple comparisons test; \**p* ≤ 0.05, \*\**p* ≤ 0.01, \*\*\*\**p* ≤ 0.0001; *n* ≥ 40 cells per cell line). (G) Graphs showing mean cell stiffness of B16F10 (left) and E0771 (right) cells treated with vehicle (Veh) or with 100 ng/ml latrunculin A (LatA) for 20 min. Data shown as mean ± SEM (two-tailed unpaired Student’s t test; *n* = 10 cells per condition), representative of 2 independent experiments. See also Fig. S6.

Although the data above are consistent with a biophysical mechanism of MRTF-induced immune vulnerability, they do not rule out the possibility that enhanced MRTF signaling might sensitize cancer cells to cytotoxic lymphocytes by changing the surface expression of a critical immunoreceptor ligand. To investigate this alternative explanation, we focused first on class I MHC and ligands for NKG2D. We were particularly interested in NKG2D because it was downregulated by human NK cells cocultured with MDA-MB-231 or MCF7 targets (Fig. S6A), implying the presence of cognate ligands on the target surface. MRTF overexpression had little to no effect on NKG2D ligands in both mouse (B16F10 and E0771) and human (MDA-MB-231 and MCF7) cell lines, and although MHC was upregulated in E0771-MRTFA cells, we did not observe increased MHC expression in B16F10-MRTFA cells or in any of the cell lines overexpressing MRTFB (Fig. S6B-C). Hence, changes in MHC or NKG2D ligands did not explain how MRTFA and MRTFB rendered cancer cells more stimulatory to cytotoxic lymphocytes.

To further investigate whether MRTF-mediated immune vulnerability was indeed caused by cytoskeletally induced biophysical changes and not the expression of undefined cell surface molecule(s), we analyzed the stimulatory capacity of giant plasma membrane vesicles (GPMVs) derived from cancer cells of interest (Fig. 7A) (Schneider et al., 2017; Sezgin et al., 2012). We reasoned that GPMVs would contain all of the molecular machinery present on the cell surface, but not the cytoskeleton, enabling us to delineate the effects of the former from the latter. Using the live F-actin probe LifeAct-GFP, we found that GPMVs did indeed lack a cortical F-actin cytoskeleton (Fig. S7A), in line with previous work (Schneider et al., 2017). Compared with whole cell extracts, GPMVs contained more of the cell surface proteins H2Kb and ATP1A1, less of the nuclear protein histone-H3, and none of the Golgi marker GM130 (Fig. S7B), consistent with a plasma membrane origin. GPMVs derived from antigen-loaded B16F10 cells stimulated CTL calcium flux and cytokine production (Fig. 6B-C), indicating that they contained the surface ligands required for T cell activation. Importantly, whereas B16F10-MRTFA and B16F10-MRTFB cells induced stronger CTL activation than control B16F10 cells (Fig. 3 and Fig. 7C), GPMVs derived from MRTF overexpressing cells were not more stimulatory than the GPMVs derived from controls (Fig. 7C), implying that the F-actin cytoskeleton is required for MRTF-induced immune activation. Using IFNγ, which drives MHC upregulation (Fig. S6B), we were able to enhance the stimulatory capacity of both B16F10 cells and the GPMVs they generated. This treatment did not, however, alter the consequences of MRTF signaling, which continued to affect only the stimulatory capacity of intact cells, but not of GPMVs (Fig. 7C). We conclude that enhanced lymphocyte stimulation by MRTF overexpressing cells is not caused by differential expression cell surface molecules, but rather by increased cytoskeletal stiffness beneath the plasma membrane.

**Fig. 7.**
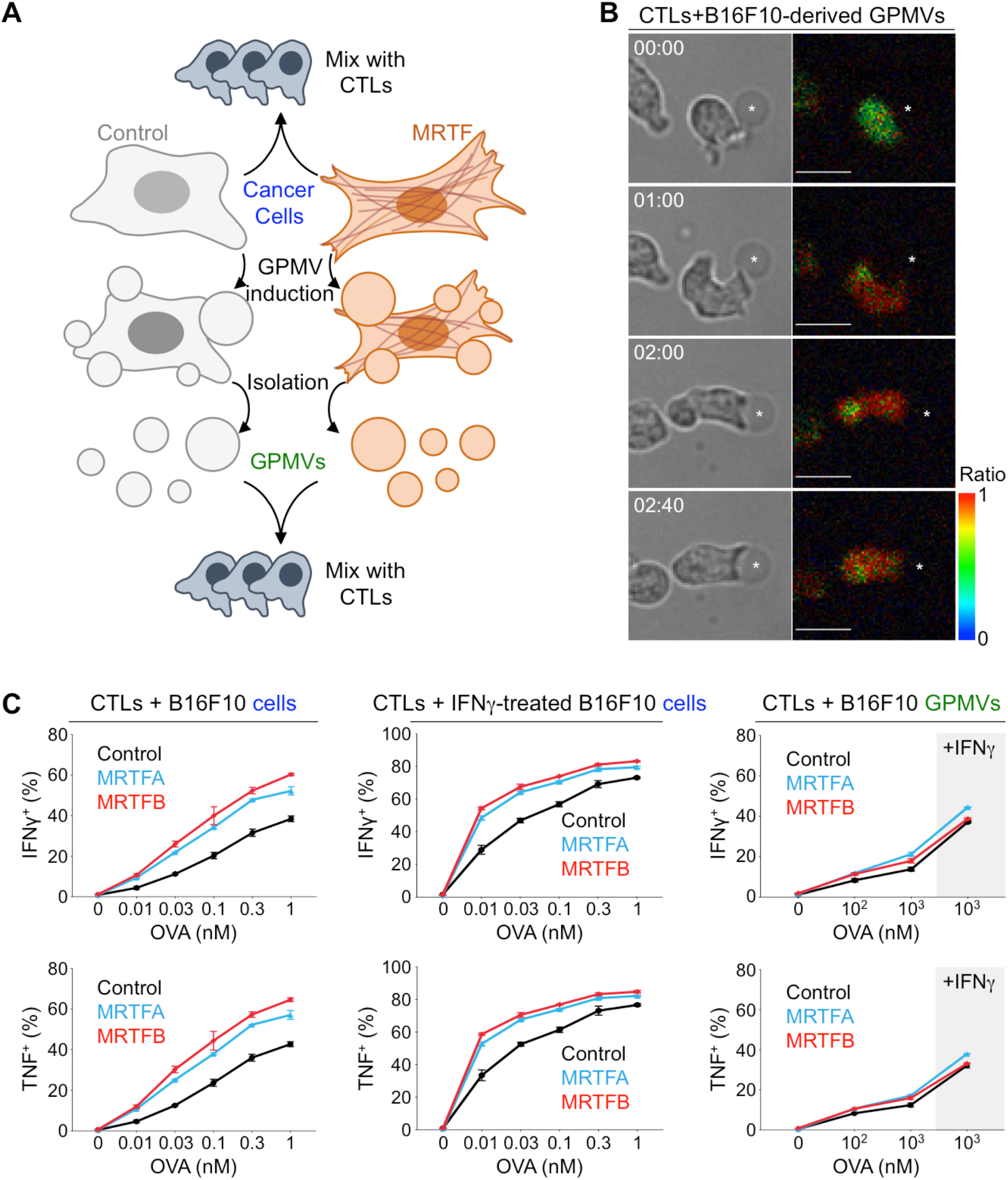
Cytoskeletal stiffness underlies the immune vulnerability of metastatic cells. CTLs were stimulated using OVA-loaded B16F10 cells or GPMVs derived from these cells. (A) Schematic diagram of experimental design (see Methods). (B) Time-lapse montage of a representative OT1 CTL Ca^2+^ response during contact with an OVA-loaded B16F10-derived GPMV (asterisk). Bright field (left) and pseudocolored Fura-2 ratio (right) are shown. Time in mm:ss is indicated in the upper left corner of each composite image. Scale bars: 10 μm. Color bar indicates the value of the Fura-2 ratio. (C) Graphs showing the percentage of CTLs producing IFN*γ* or TNF, measured by intracellular antibody staining 4 h after mixing with B16F10 cells (left), B16F10 cells pretreated with IFNγ (center), or GPMVs derived from untreated or IFNγ-treated B16F10 cells (right). Data shown as mean ± SEM of technical triplicates, representative of 3 independent experiments. See also Fig. S7.

## DISCUSSION

Characteristic genetic and biochemical traits enable cancer cells to grow in an unregulated manner, but they also create vulnerabilities, such as oncogene addiction, metabolic reliance, and sensitivity to genotoxic agents, that can be targeted by appropriate therapeutic modalities (Behan et al., 2019; Bryant et al., 2005; DeBerardinis and Chandel, 2016; Farmer et al., 2005; Weinstein, 2002). Our present study extends this paradigm into the biophysical domain by demonstrating that the architectural and mechanical properties that enable metastatic growth also serve as an Achilles heel for destruction by cytotoxic lymphocytes. Isolated cancer cells are typically less rigid than untransformed cells from the same parent tissue (Guck et al., 2005; Hou et al., 2009; Xu et al., 2012). To occupy the metastatic niche, however, cancer cells must spread on the microvascular basement membrane (Er et al., 2018; Valiente et al., 2014), thereby increasing their rigidity to the point where they trigger robust lymphocyte activation. This coupling of colonization with biophysical vulnerability provides an explanation for why metastasis is so inefficient, and it also identifies mechanosensing of cancer cell rigidity as a novel mode of immunosurveillance.

In principle, mechanical immunosurveillance, or mechanosurveillance, would enable the immune system to target cellular dysfunction that does not detectably alter the biochemical recognition of cell surface proteins or secreted factors. In practice, however, it seems more likely that both biochemical and biophysical features will control immune vulnerability in a combinatorial manner. For instance, our observation that MRTF-induced suppression of E0771 cells requires CD8^+^ CTLs, while the anti-B16F10 response is dominated by NK cells, probably reflects the fact that E0771 cells express higher levels of class I MHC, which would activate T cells and inhibit NK cells. Hence, the cell biological contexts within which mechanosurveillance operates will be dictated by specific molecular interactions between cytotoxic lymphocytes and the target cells in question. Deciphering this crosstalk will be a fascinating area of future study.

It is generally thought that increasing target rigidity amplifies lymphocyte activation via mechanosensitive cell surface receptors (Huse, 2017; Zhu et al., 2019). Certain immunoreceptors, including the TCR, integrins, and the NK receptor CD16, are known to form catch bonds, in which the lifetime of interactions with cognate ligand increases under applied force (Gonzalez et al., 2019; Kong et al., 2009; Liu et al., 2014). There are also indications that synaptic forces induce conformational changes in the TCR, integrins, and components of integrin-mediated adhesions that are required for optimal signal transduction (del Rio et al., 2009; Friedland et al., 2009; Lee et al., 2015). Actin dependent stiffening of the target cell could facilitate all of these processes by restraining deformation orthogonal to the cell surface. Enhanced cortical F-actin accumulation could also restrict the lateral mobility of cell surface ligands in the plasma membrane by strengthening adhesion between the membrane and the cytoskeleton or by altering the confinement properties of membrane corrals (Gauthier et al., 2012; Jacobson et al., 2019). Reduced lateral mobility is a particularly interesting possibility in light of work indicating that dendritic cells potentiate integrin activation on T cells by restraining the diffusion of an integrin ligand on their own surface (Comrie et al., 2015).

Although cortical F-actin is a predominant regulator of cellular mechanics, other molecular components influence the biophysical properties of cells and could therefore contribute to mechanosurveillance. Ezrin-radixin-moesin (ERM) proteins and lipid modifying enzymes modulate cellular architecture and migration by controlling interactions between the plasma membrane and the F-actin cortex (Balla, 2013; Clucas and Valderrama, 2014). Both classes of protein have been implicated in cancer progression and therefore represent intriguing candidate regulators. Oncogenic transformation is also associated with dysregulation of the microtubule cytoskeleton (Parker et al., 2014), which could result in structural abnormalities that are detectable by immune cells. A role for microtubules in mechanosurveillance is particularly intriguing because they are targeted by a number of chemotherapeutic agents, such as Taxol, raising the possibility that these treatment modalities might modulate anti-tumor immune responses biophysically.

The immune sensitization mechanism characterized in this study resulted from a cell intrinsic mechanical trait (cellular stiffness). That being said, tumor progression is associated with cell extrinsic biophysical changes, as well, such as increased ECM adhesiveness and rigidity (stromal stiffness) (Kai et al., 2019; Levental et al., 2009), which could also be coupled to immune vulnerabilities. ECM remodeling is generally thought to promote malignancy by stimulating cancer cell migration, epithelial to mesenchymal transition (EMT), and transcriptional programs important for tumorigenesis (Kai et al., 2019). ECM stiffness, in particular, has been shown to drive cancer cell proliferation via the transcription factor YAP (Yes-associated protein) (Albrengues et al., 2018; Panciera et al., 2020). These same changes in the ECM, however, could also promote anti-tumor immunity in certain contexts. Indeed, matrix-induced metastatic outgrowth drives MRTF signaling (Er et al., 2018), which we now know triggers lymphocyte mechanosurveillance. Further study of how the ECM and other aspects of tumor architecture and mechanics affect both the movement and the functional potential of infiltrating immune cells could identify additional therapeutic opportunities.

The idea that metastatic cells must fine-tune MRTF signaling to balance the benefits of cell spreading with the drawbacks of immune activation conceptually parallels recent work on the regulation of EMT during cancer progression (Alderton, 2013; Ocana et al., 2012; Tsai et al., 2012). Although EMT promotes migration, invasion, multipotency, and resistance to certain therapies (Zhang and Weinberg, 2018), it has also been shown to hinder proliferation and promote apoptosis (David et al., 2016; Kong et al., 2017). Hence, to metastasize effectively, cancer cells must employ transcriptional programs of both EMT and mesenchymal to epithelial transition, balancing the proliferative and invasive properties of each cellular state. The importance of fine tuning EMT in this way is highlighted by the observation of partial EMT signatures in patients with metastatic disease (Puram et al., 2018). Interestingly, the loss of some, but not all, epithelial characteristics during partial EMT is thought to be important for the acquisition of stem cell-like properties and metastatic dormancy, a state of prolonged quiescence in which cancer cells evade the immune system (Lawson et al., 2015; Malladi et al., 2016; Pommier et al., 2018). Dormancy ends when metastatic colonies engage the ECM, thereby activating YAP and MRTF to drive outgrowth (Albrengues et al., 2018; Er et al., 2018; Shibue and Weinberg, 2009). Determining how these various cellular states affect not only the biochemical but also the biophysical properties of cancer cells will provide for a better understanding of how metastatic tumors balance partial EMT, stem cell-like behavior, and awakening from metastatic dormancy in order to grow in the face of immunosurveillance.

The RAC1^P29S/L^ allele is generally associated with enhanced melanoma malignancy and resistance to targeted therapies (Lionarons et al., 2019; Van Allen et al., 2014; Watson et al., 2014), including the BRAF inhibitor vemurafenib. Our results, however, indicate that this mutation actually increases tumor responsiveness to ICB, implying that RAC1^P29S/L^ may be useful as a positive predictive indicator for this class of treatments. Although RAC1 activates multiple downstream signaling pathways, MRTF has been shown to be critical for the specific effects of RAC1^P29S^ on melanoma physiology (Lionarons et al., 2019), and our mechanistic results *in vitro* and in mouse models support a role for MRTF in triggering anti-tumor immunity, specifically via mechanosurveillance. Critically, we have also documented an MRTF-SRF gene signature in human RAC1/2^P29S/L^ melanoma, providing direct evidence that MRTF signaling enhances ICB-induced immunosurveillance of human tumors. It will be interesting to see if the MRTF-SRF signature itself correlates with responsiveness to ICB, independent of RAC1 mutation, as this would enable identification of additional patients likely to benefit from this therapy.

A number of intracellular pathogens, including HIV, Chlamydia, and Listeria, dramatically remodel the host cell cytoskeleton to enable intracellular motility, proliferation, and the infection of neighboring cells (Bhavsar et al., 2007; Metais et al., 2018; Wesolowski and Paumet, 2017). The capacity of these architectural changes to modulate the immune response, both positively and negatively, has not been examined. It has been shown, however, that CD4^+^ T cells latently infected with HIV contain high levels of viral integration in the *Mrtfa* and *Mrtfb* loci, implying that disruption of the MRTF pathway enables infected cells to elude the immune system (Maldarelli et al., 2014). Furthermore, the HIV virulence factor Nef, which disrupts cytoskeletal polarity, was recently found to enhance the survival of infected cells in an immunocompetent mouse model (Usmani et al., 2019). Therefore, defining the genetic and molecular bases of mechanosurveillance will likely illuminate cellular immunity against not only cancer but also infectious disease.

## METHODS

### Constructs

Retroviral vectors for LifeAct-GFP, GFP, and TGL expression have been described(Le Floc’h et al., 2013; Ponomarev et al., 2004). Constructs for MRTFA/B silencing and overexpression were gifts from Ron Prywes (Addgene # 27161, #19846, and #27175). To generate inducible expression constructs, the coding sequences of MRTFA and MRTFB were PCR amplified and subcloned into the pRetroX-Tight-Hygro vector (Takara Bio, 631034). Doxycycline inducible expression was achieved by pLVX-TetON-Advanced (Takara Bio, 632162) expression.

### Mice and cell culture

The animal protocols used for this study were approved by the Institutional Animal Care and Use Committee of Memorial Sloan Kettering Cancer Center. Recipient C57BL/6J mice for *in vivo* assays were purchased from the Jackson Laboratory. To generate OT1 CTLs, T cells from OT1 αβTCR transgenic mice (Taconic) were mixed with congenic splenocytes pulsed with 100 nM OVA and cultured in RPMI medium containing 10 % (vol/vol) FBS. Cells were supplemented with interleukin 2 (IL-2, 30 IU/ml, NIH BRB Repository) after 24 h and were split as needed in RPMI containing IL-2 and used for functional assays after 7 days in culture. Murine NK cells were isolated from C57BL/6J splenocytes by negative selection using an NK cell isolation kit (MACS, 130-115-818) and incubated overnight in 1000 U/ml IL-2. Human NK cells were isolated from peripheral blood samples obtained from healthy volunteer donors via the New York Blood Center (NYBC, http://nybloodcenter.org/). The Memorial Sloan Kettering Cancer Center Institutional Review Board (MSKCC IRB) waived the need for additional research consent for anonymous NYBC samples. Peripheral blood mononuclear cells (PBMCs) were purified from buffy coats by density gradient centrifugation (Ficoll-Paque Plus; GE Healthcare) and then cryopreserved in FBS with 10% DMSO. One day prior to the experiment, cells were thawed and incubated in clone media (DMEM, 30% Ham’s F-12, 10% human serum, 1 mM sodium pyruvate, 1% MEM nonessential amino acids, 2 mM L-glutamine, 50 U/ml penicillin, 50 μg/ml streptomycin) supplemented with 200 U/ml IL-2 (Proleukin, Prometheus) at 37 °C. B16F10 and E0771 cell lines were cultured in RPMI, while MCF7 and MDA-MB-231 cell lines were cultured in DMEM. Media were supplemented with 10 % FBS, 1 mM sodium pyruvate, 2 mM L-glutamine, 50 U/ml penicillin, and 50 μg/ml streptomycin. All MDA-MB-231 experiments utilized the brain metastatic subline MDA-231Br (Bos et al., 2009). Cell lines to assess the effects of inducible MRTFA/B overexpression were prepared by sequential transduction with rTTA, TGL, and either control or MRTF overexpression vectors, followed by culture in G418 and hygromycin. MRTF expression was induced by treating cells with 500 ng/ml doxycycline hyclate (Sigma) 24-48 h prior to the experiment.

### Metastasis assays

2 × 10^5^ B16F10 or E0771 cells were injected into the tail vein of 4-6 week old C57BL/6J mice (Jackson Labs, 000664). Doxycycline was included in the mouse food (2500 mg/kg) to maintain MRTF expression for the duration of the experiment. Mouse hair was removed using clippers to prevent interference with bioluminescent imaging (BLI). Lung metastasis burden was quantified weekly using retro-orbital D-luciferin (150 mg kg−1) injection followed by imaging via the IVIS Spectrum Xenogen instrument (Caliper Life Sciences) installed with Living Image software v.2.50. Metastatic load per mouse was calculated by dividing the total photon flux signal at the end point of the experiment by the total photon flux measured immediately after cancer cell delivery on the day of injection. NK cell depletion was performed by injecting mice intraperitoneally (i.p.) with anti-asialo GM1 antibody (Wako Chemicals, 986-10001) as previously described (Er et al., 2018) 6 days and 1 day before tail vein injection of cancer cells and once a week thereafter. CD8 positive T cell depletion was achieved using 250 μg InVivoMab anti-mouse CD8α antibody (clone 53-6.7, BioXCell, BE0004-1) or IgG2a control (BioXCell, BE0089), injected 2 days and 1 day before tumor delivery and once every week thereafter. Immune checkpoint blockade was achieved by injecting mice i.p. with 125 μg anti-CTLA-4 (clone 9D9) or mouse control IgG2b (clone MPC-11) antibodies 4 days after delivery of cancer cells.

### Killing, lytic granule secretion, and cytokine production assays

For the CTL functional assays, cancer cell targets were cultured overnight on fibronectin-coated 96-well plates. They were then loaded with varying concentrations of OVA for 2 h and washed three times in medium. To assess killing, OT1 CTLs were added at a 4:1 effector to target (E:T) ratio and incubated for 5 h at 37 °C in culture medium. Cells were then labeled with APC conjugated anti-CD8a antibody (Tonbo Biosciences, 20-0081), and specific lysis of target cells (GFP^+^, CD8^−^) was determined by propidium iodide (PI, Thermo Fisher Scientific) incorporation using flow cytometry. To assess lytic granule secretion, the E:T ratio was 2:1, and cells were incubated for 90 min at 37 °C in the presence of eFluor660 conjugated anti-Lamp1 antibody (1 μg/ml, Clone 1D4B, eBiosciences). Cells were then labeled with anti-CD8a antibody, and the percentage of CTLs (CD8^+^) with positive Lamp1 staining was quantified by flow cytometry. To assess cytokine production, the E:T ratio was 2:1, and cells were incubated for 4 h at 37 °C in the presence of BD GolgiPlug™ protein transport inhibitor (BD Biosciences). Cells were then labeled with anti-CD8a antibody and a dead cell marker (Live/Dead Fixable Aqua Dead Cell Stain Kit), fixed, and permeabilized using the BD Cytofix/Cytoperm™ kit. After labeling with PE conjugated anti-TNF (BioLegend, 506306) and PE/Cy7 conjugated anti-IFNγ (BioLegend, 505826) antibodies, the percentage of CTLs (CD8^+^) expressing TNF and IFNγ was determined by flow cytometry. All functional assays were performed in triplicate. For mouse NK cell functional assays, cancer cell targets were cultured overnight on fibronectin-coated 96-well plates. They were then mixed with NK cells at a 1:1 ratio and incubated for 6 h at 37 °C in the presence of eFluor660 conjugated anti-Lamp1 antibody. Subsequently, cells were labeled with PerCP-Cy5.5 conjugated anti-NK1.1 antibody (eBioscience, 45-5941-82) and the percentage of NK1.1^+^ cells with positive Lamp1 staining was quantified by flow cytometry. For human NK cell functional assays, cancer cell targets were cultured overnight on fibronectin-coated 96-well plates. To assess NK cell degranulation, 2 × 10^5^ PBMCs were added to each well (NK cells comprise 5-15 % of PBMCs) in the presence of monensin (GolgiSTOP™; 1:1,000 dilution; BD) and Brilliant Violet 786-labeled anti-Lamp1 mAb (clone SJ25C1, BD Horizon) for 5 h at 37 °C. After incubation, cells were collected in a 96-well V-bottom plate, washed and stained with dead cell marker (Live/Dead Fixable Near IR Dead Cell Stain Kit), ECD-labeled anti-CD56 mAb (Beckman Coulter), BV650-labeled anti-CD3 mAb (clone UCHT1, BD Horizon), and PE-labeled anti-NKG2D (clone 1D11, BioLegend). Finally, cells were washed in 1% FBS/PBS and subjected to flow cytometry (LSR Fortessa). All flow cytometric analysis was performed using FlowJo software.

### *In vitro* cell growth and proliferation assays

To assess cell viability, the CellTiter-Glo® Luminescent Cell Viability Assay Kit (Promega, G7570) was used according to manufacturer instructions. To assess cell proliferation, cells were labeled with CellTrace Violet (CTV, Thermo Fisher) according to manufacturer instructions, and CTV dilution was quantified by flow cytometry.

### Cell death assays

Cell death was induced by treating cells seeded on fibronectin-coated 96 well plates with varying concentrations of staurosporine (Cell Signaling Technology), FasL (PeproTech), TNF (PeproTech), and granzyme B (BioLegend). Granzyme B was activated using Cathepsin C/DPPI (R&D Systems) according to manufacturer instructions, and applied to cells in combination with a sublytic concentration of perforin, which was purified as previously described (Basu et al., 2016). Cell death after 5 h treatment was quantified by PI incorporation or using the Caspase Glo system (Promega) according to manufacturer instructions.

### Transcriptome sequencing and analysis

RNA was collected using the RNeasy Mini Kit (Qiagen, 74106) according to manufacturer instructions. After RiboGreen quantification and quality control by Agilent BioAnalyzer, 500 ng of total RNA underwent polyA selection and TruSeq library preparation according to instructions provided by Illumina (TruSeq Stranded mRNA LT Kit), with 8 cycles of PCR. Samples were barcoded and run on a HiSeq 4000 in a 50bp/50bp paired end run, using the HiSeq 3000/4000 SBS Kit (Illumina). An average of 41 million paired reads was generated per sample and the average fraction of mRNA bases was 74%. Output data were mapped to the target genome with the rnaStar aligner (Dobin et al., 2013) using the 2 pass mapping method (Engstrom et al., 2013). After postprocessing with PICARD, the expression count matrix was computed using HTSeq (www-huber.embl.de/users/anders/HTSeq). The raw count matrix generated by HTSeq was then processed in DESeq (www-huber.embl.de/users/anders/DESeq) to normalize the full dataset and analyze differential expression between sample groups. Gene Ontology analysis of Biological Processes and Cellular Components was performed using the DAVID 6.8 Functional Annotation Tool (Huang da et al., 2009) with Benjamini correction for multiple hypothesis testing and a cut-off of 2 genes minimum per cluster. Each Gene Ontology table in Fig. 5 and Fig. S5 lists the 10 annotation clusters with the highest enrichment scores and lowest *p* values below *p* ≤ 0.05.

### Analysis of clinical data

Patient analysis was carried out using cBioportal (Cerami et al., 2012; Gao et al., 2013). The TCGA PanCancer Atlas database (Ellrott et al., 2018; Gao et al., 2018; Hoadley et al., 2018; Liu et al., 2018; Sanchez-Vega et al., 2018; Taylor et al., 2018) was used for survival analysis of RAC1/2^P29S/L^ skin cutaneous melanoma patients (Supplementary Table 1). Gene set enrichment analysis (GSEA) of SRF target gene expression enrichment was performed using data from TCGA skin cutaneous melanoma patients that were positive or negative for the following mutations: RAC1/2^P29S/L^, NF1 truncation mutant, PPP6C^R264C^, FBXW7 mutant, and IDH1^R132C/L^. RNA-sequencing read count estimation values using RNA-Seq by Expectation Maximization (RSEM) were downloaded from the Genomics Data Commons Data Portal. The SRF_01 gene set (M12047) from the Molecular Signatures Database (MSigDB v7.0) was used for GSEA analysis. For heatmap analysis, RAC1/2^P29S/L^(+) and RAC1/2^P29S/L^(−) samples were grouped and clustered using leading genes in the SRF_01 gene set GSEA. For survival analysis of RAC1/2^P29S/L^, NF1 truncation mutant, PPP6C^R264C^, FBXW7 mutant, and IDH1^R132C/L^ patients treated with anti-CTLA4, patients that underwent anti-CTLA4 therapy were selected from reference studies (Catalanotti et al., 2017; Liang et al., 2017; Miao et al., 2018; Samstein et al., 2019) (Supplemental Table S2).

### Image analysis

To quantify F-actin intensity in fixed cells, each image was subjected to intensity thresholding in Imaris (Bitplane) to establish the space occupied by cells, after which the average intensity of Alexa Fluor 594-labeled phalloidin within the cellular volume was determined. To quantify *in vivo* proliferation, mouse lungs bearing metastatic tumors were paraffin embedded, sectioned into 5 μm thick slices, stained for Ki67 (Cell Signaling Technologies, 9129) together with DAPI, and imaged using a 3DHISTECH Pannoramic Scanner with a 20 × objective lens. The number of cell nuclei (DAPI) and proliferating cells (Ki67) were counted using automated segmentation. Code available upon request.

### Cell surface proteins

Target cells were gently detached using Trypsin/EDTA and washed prior to antibody staining. B16F10 and E0771 cells were stained with APC-labeled anti-Fas (clone SA367H8, BioLegend), PE-labeled anti-H2Kb (clone AF6-88.5, BioLegend), PE-labeled anti-H2Db (clone 28-14-8, eBioscience), and mouse NKG2D-Fc (kind gift from J. C. Sun) followed by a PE-labeled secondary antibody. MDA-MB-231 and MCF7 cells were stained with PE-labeled anti-MICA/B (clone 6D4, BD Pharmingen) and PE-Cy5-labeled anti-HLA-ABC (G46-2.6, BD Pharmingen). Surface expression was then quantified by flow cytometry.

### Atomic Force Microscopy (AFM)

Cells were seeded on glass-bottom petri dishes (FluoroDish FD5040) coated with fibronectin (from bovine plasma, Millipore Sigma) and then kept in complete RPMI medium with 10 mM HEPES pH 7.0 during the acquisition of stiffness maps. Experiments were performed at 37 °C with an MFP-3D-BIO AFM microscope (Oxford Instruments) using cantilevers with 5 μm diameter colloidal borosilicate probes (nominal spring constant k = 0.1 N/m, Novascan). Before each experiment, the exact spring constant of the cantilever was determined using the thermal noise method and its optical sensitivity determined using a PBS-filled glass bottom petri dish as an infinitely stiff surface. 10-12 cells from each experimental group were tested in each session. Bright field images of each cell were collected during AFM measurements using an inverted optical objective (Zeiss AxioObserver Z1) integrated with the AFM. Stiffness maps of 60 × 60 μm^2^ (18 × 18 points) were collected in areas containing both cells and substrate at 1.5 Hz for a single approach/withdraw cycle. A trigger point of 1 nN was used to ensure sample penetration of 1-2 μm. Force curves in each map were fitted according to the Hertz model (Igor Pro, Wavemetrics). Data fitting was performed in the range from 0 to 50% of the maximum applied force to consider only measurements within the first 1 μm of indentation. The following settings were used: tip Poisson ν_tip_ = 0.19, tip Young’s modulus E_tip_ = 68 GPa, and sample Poisson ν_sample_ = 0.45. Stiffness histograms were obtained by identifying the stiffness values belonging to each cell (and not the substrate values, shown in black on force maps) through a mask and plotting the results from each cell line as a single population. All measurements made < 500 nm above the substrate were excluded. Extraction of the stiffness from the raw Igor Binary Wave (.ibw) data with an overlapped mask was obtained by means of a home-built routine implemented in Igor (Igor Pro, Wavemetrics). Stiffness distribution histograms were obtained using the histogram analysis tool in Excel (Microsoft) after normalizing for the total number of data points.

### Giant plasma membrane vesicle (GPMV) isolation and purification

GPMVs were generated as previously described (Sezgin et al., 2012) with minor modifications for scale and cell type. 1.5 × 10^6^ cells were seeded in a 10 cm^2^ dish and incubated for 18 h with or without 25 ng/ml IFN*γ* and the indicated concentrations of OVA peptide. Cells were then transferred into 5 ml of GPMV buffer (10 mM HEPES, 150 mM NaCl, 2 mM CaCl_2_, 25 mM PFA, 2 mM DTT, pH 7.4) for 1 h at 37 °C. To purify GPMVs, the suspension was centrifuged at 100 × g for 10 min to pellet cell debris, and then the supernatant was centrifuged at 2000 × g for 1 h at 4 °C to pellet the GPMVs. For lymphocyte stimulation, GPMVs were washed and then resuspended in 500 μl of cell culture medium. 100 μl of this purified GPMV sample was then mixed with 10^4^ CTLs in a v-bottom 96-well plate and incubated for the indicated times. For immunoblot analysis of GPMV protein content, GPMV pellets were resuspended in 1 × Alfa Aesar Laemmli SDS Sample Buffer (Fisher Scientific, AAJ61337AC) prior to gel electrophoresis. For imaging studies (Fig. S7A), B16F10 cells transiently transfected with GFP or LifeAct-GFP were labeled with CellMask™ Orange Plasma Membrane Stain according to manufacturer instructions, and then either imaged or used to generate GPMVs for imaging.

### Ca^2+^ imaging of CTLs and GPMVs

CTLs were loaded with 5 μg/ml Fura-2AM and then added to poly-L-lysine-coated chambers containing GPMVs derived from OVA-loaded B16F10 cells. Fura-2 images using 340 nm and 380 nm excitation were acquired every 30 seconds for 20 min, using a 20 × objective lens fitted to an IX-81 microscope stage (Olympus).

### Statistics

Analyses were carried out using either representative experiments or pooled data as indicated (*n* is defined in the figure legends for each experiment). Statistical tests (two-tailed Mann-Whitney, two-tailed ANOVA, paired and unpaired two-tailed t tests and Log-rank Mantel-Cox tests) were performed using GraphPad Prism. Log-rank tests for patient survival were implemented in cBioportal (Cerami et al., 2012; Gao et al., 2013). Unless otherwise indicated, error bars denote SEM. No statistical methods were used to determine sample size prior to experiments.

---

**Fig. S1.**
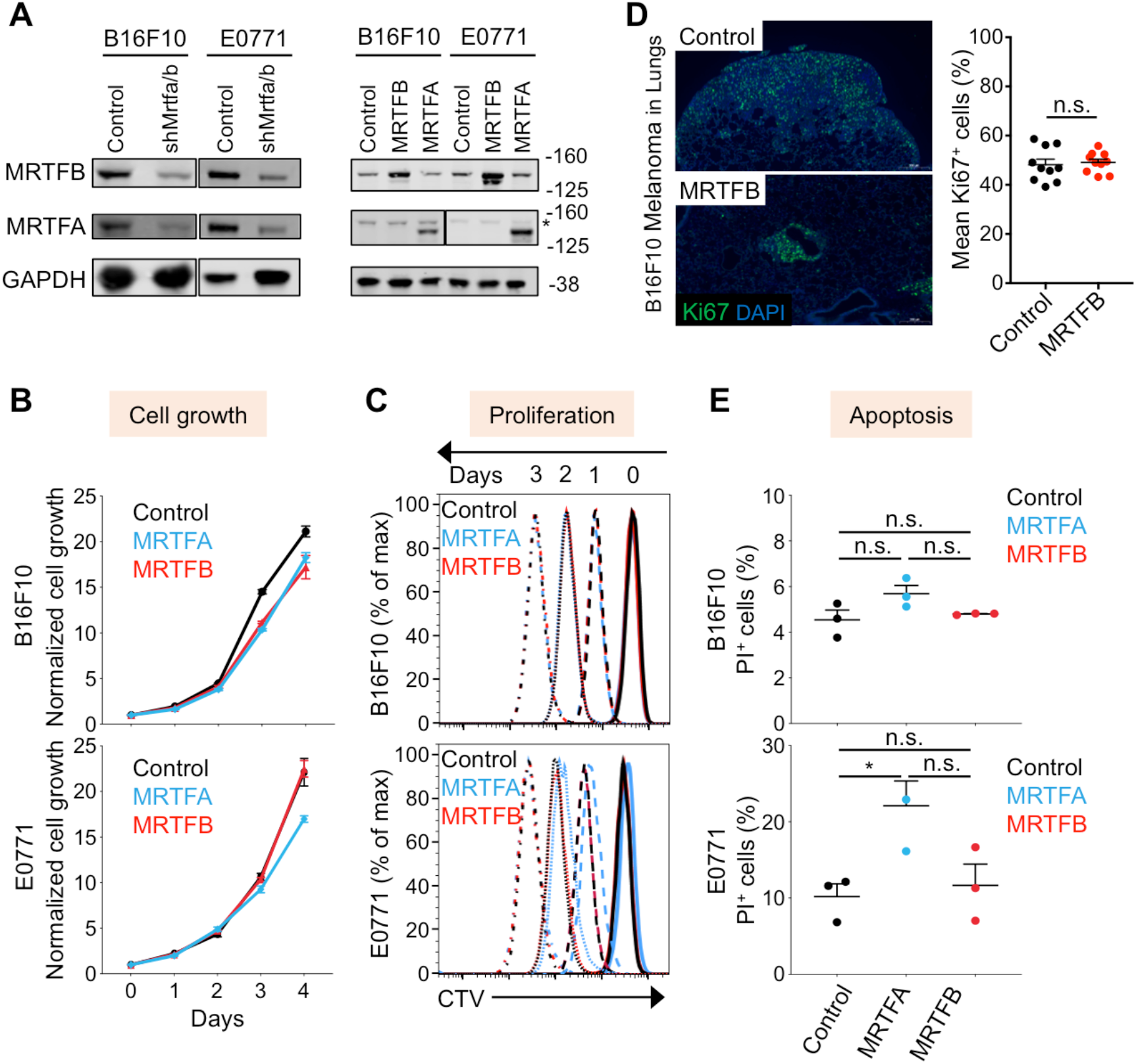
MRTF does not affect cancer cell growth, proliferation or apoptosis. Related to Fig. 1 and 2. (A) Representative western blots showing MRTFA/B expression levels in the indicated B16F10 and E0771 cell lines. * indicates endogenous MRTFA. (B) Cellular growth kinetics of the indicated B16F10 (above) and E0771 (below) cell lines, measured by CellTiterGlo normalized to first day of plating (Day 0). Error bars: SEM. *n* = 3 technical replicates. Results are representative of two independent experiments. (C) Cellular proliferation measured by CTV (Cell Trace Violet) dilution, after CTV staining on day 0. Data are representative of 3 independent experiments. (D) Left, immunofluorescence images showing proliferative state (Ki67, green) of B16F10 melanoma cells expressing control vector or MRTFB during lung colonization of syngeneic mice. DAPI, nuclear stain, blue. Right, quantification of Ki67 staining, with data shown as mean ± SEM, n.s.: not significant for *p* > 0.05; two-tailed unpaired Student’s t test (*n* = 10 mice per group). (E) Graphs showing the percentage of cell apoptosis quantified by propidium iodide (PI) incorporation. Data from 3 independent experiments shown as mean ± SEM (One-way ANOVA with Tukey’s multiple comparisons test; n.s.: not significant for *p* > 0.05, \**p* ≤ 0.05).

**Fig. S2.**
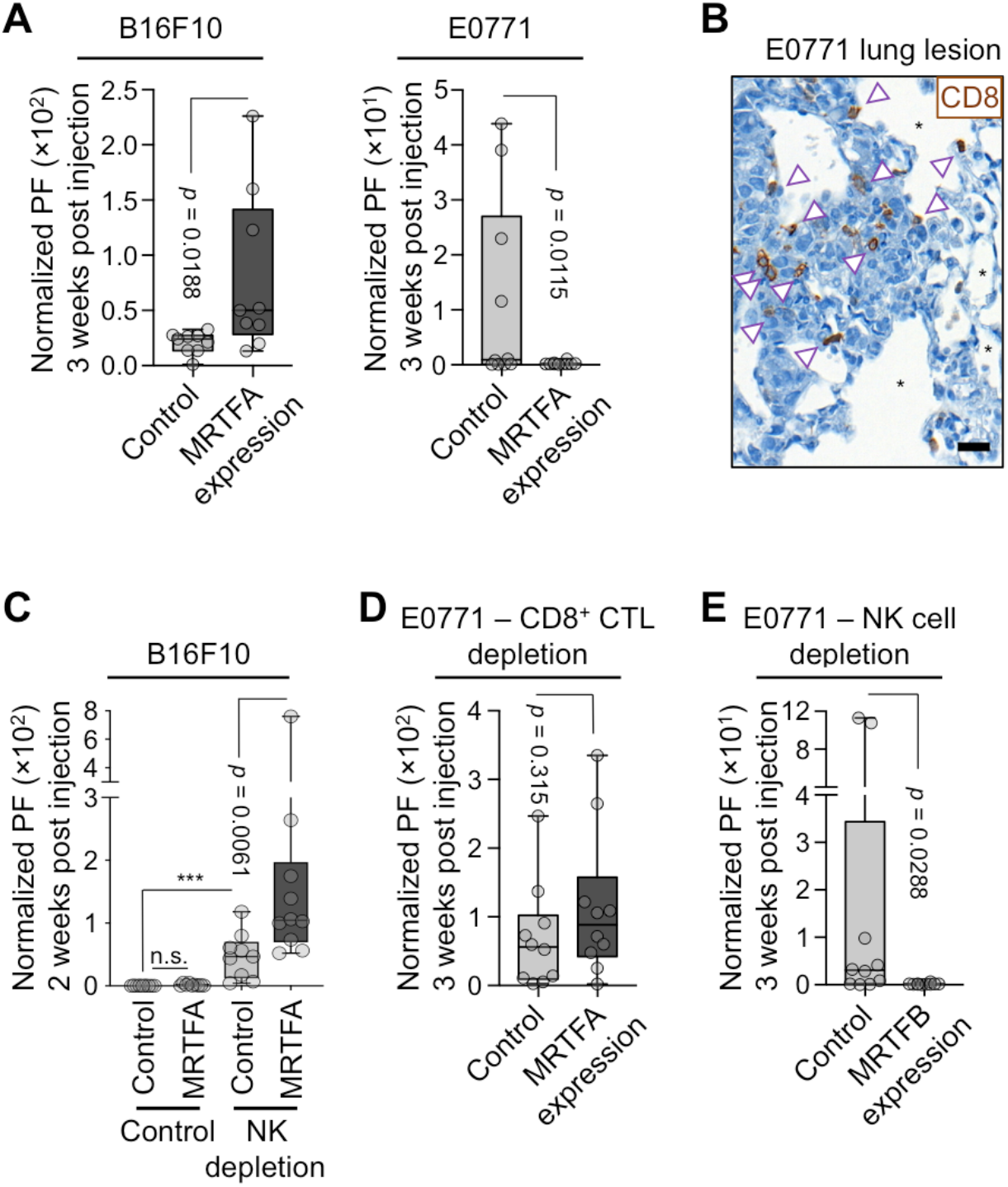
Immune vulnerability of cancer cells during metastatic colonization. Related to Fig. 1 and 2. (A) BLI of mouse lungs 3 weeks post tail vein injection with B16F10 melanoma (left) or E0771 breast cancer (right) cells overexpressing empty vector or MRTFA (*n* = 9 mice per group). (B) Representative IHC images of CD8^+^ T cell (brown arrowheads, CD8 staining) infiltration in E0771 breast cancer lung metastases. *: alveolar space, Scale bar: 20 μm. (C) BLI of mouse lungs 2 weeks after tail vein injection with B16F10 cells with or without NK cell depletion using anti-Asialo GM1 antibody, showing sensitivity of MRTFA expressing cells to NK cells during lung colonization. n.s. :not significant, *p* = 0.077, \*\*\*\**p* < 0.0001 (*n* = 10 mice for MRTFA, NK cell depletion and *n* = 9 mice for others). (D-E) BLI of mice pretreated with anti-CD8 antibody (D) or anti-asialo GM1 antibody (E) for T and NK cell depletion, respectively, and imaged 3 weeks after injection of indicated cancer cells (*n* = 10 mice per group). *p* values were calculated by Mann-Whitney test.

**Fig. S3.**
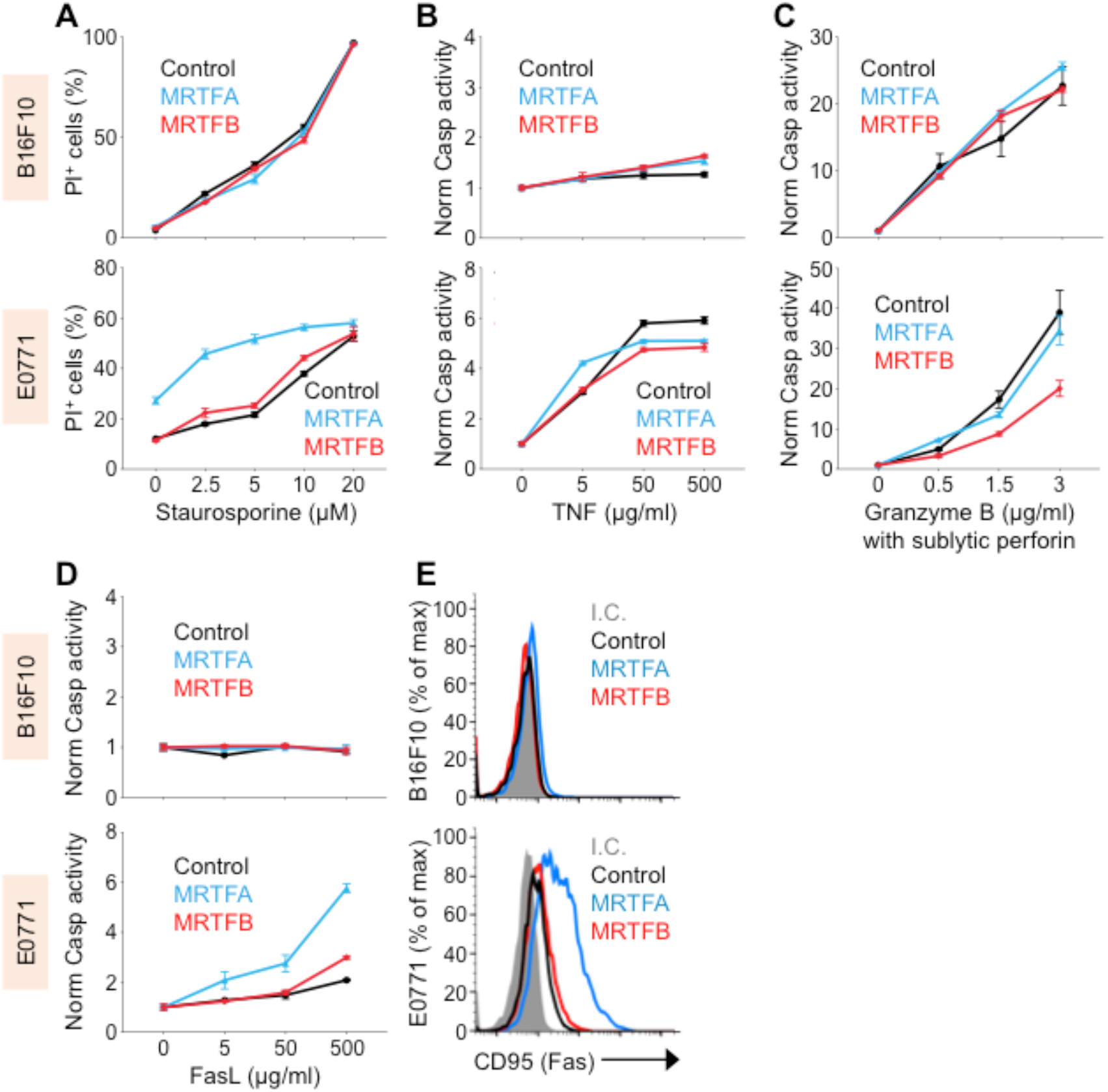
MRTF does not increase cancer cell sensitivity to apoptotic insults. Related to Fig. 3. (A-D) B16F10 (top) and E0771 (bottom) control and MRTFA or MRTFB overexpressing cell lines were treated with the indicated concentrations of staurosporine (A), TNF (B), granzyme B plus a sublytic concentration of perforin (C), or FasL (D) and cell apoptosis was quantified by propidium iodide (PI) incorporation (A) or the caspase Glo 3/7 assay system after normalization to each cell line’s untreated samples (B-D). All data shown as mean ± SEM of technical triplicates, representative of 2 independent experiments. (E) Flow cytometric analysis of Fas on the indicated control and MRTFA/B overexpressing cell lines. Isotype control (I.C.) is shown in gray. Histograms representative of 3 independent experiments.

**Fig. S4.**
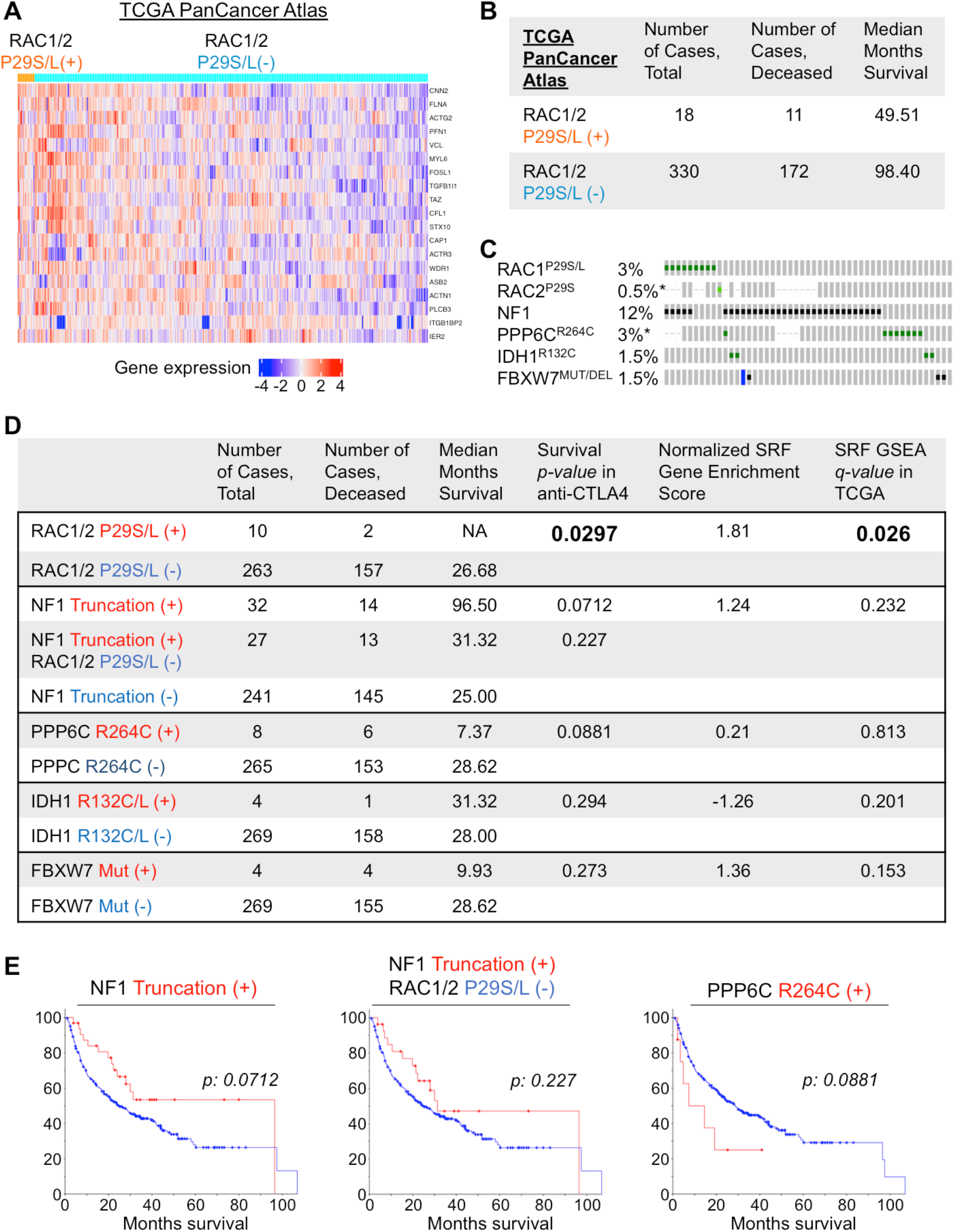
An MRTF-SRF gene signature correlates with responsiveness to ICB. Related to Fig. 4. (A) Heat map representation of differential gene expression z-scores for the leading genes in the MRTF-SRF gene signature applied to the TCGA PanCancer Atlas skin cutaneous melanoma dataset. (B) Total number of patients, their RAC1/2 mutational and vital status, and median patient survival in months for the TCGA skin cutaneous melanoma data set. (C) Oncoprint showing the overlap between distinct sets of mutations in patients from the anti-CTLA4 cohort analyzed in Fig. 4H. Each grey bar is a profiled patient, with dashes representing no available data on a particular gene. Color-coding on each bar represents a mutation or a genomic alteration annotated as oncogenic by oncoKB. Green and black squares are oncogenic driver point mutations and truncations, respectively. Blue bars denote deep genomic deletions. n = 273 patients. * indicates that not all patients were profiled for the queried genomic event. Patients with no alterations were included in the analyses but were cropped from the panel for simplicity. (D) Survival statistics for mutations depicted in C. The first four columns contain statistics from the anti-CTLA4 cohort, while the last two columns of SRF signature statistics are derived from the TCGA skin cutaneous melanoma cohort. Bolded values are statistically significant. (E) Kaplan-Meier curves showing overall survival of melanoma patients from the anti-CTLA4 cohort bearing the indicated UV-induced mutations. *p* values were calculated by Log-rank test.

**Fig. S5.**
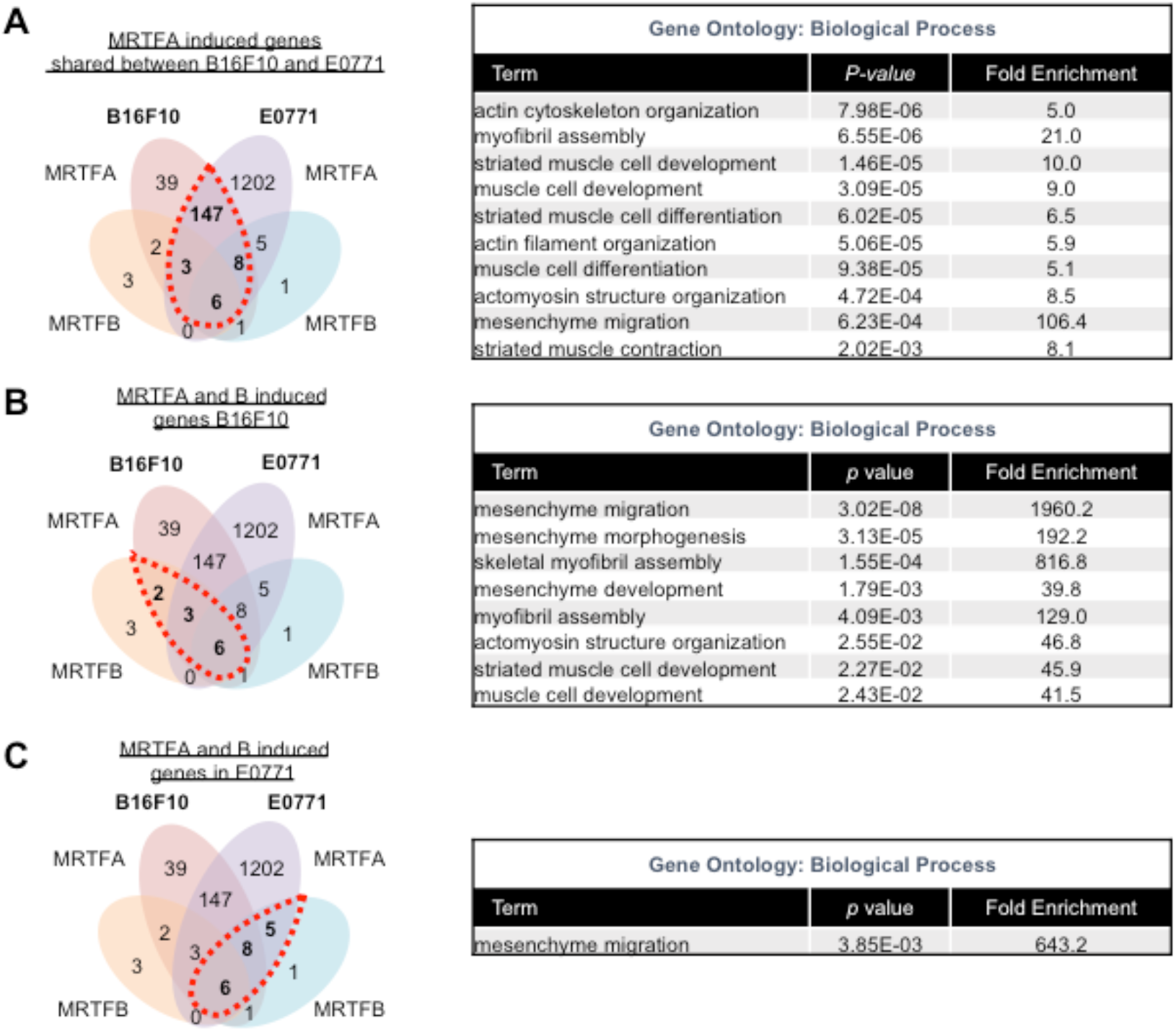
RNA sequencing analysis of MRTFA and MRTFB expressing cells. Related to Fig. 5. (A-C) Left, Venn diagrams of upregulated genes exclusive to or shared by B16F10 and E0771 cells, with bold numbers and red dashed circles highlighting gene sets used for the GO analyses on the right. Up to 10 statistically significant GO terms are shown in each table, with reported *p* values corrected for multiple testing using the Benjamini method.

**Fig. S6.**
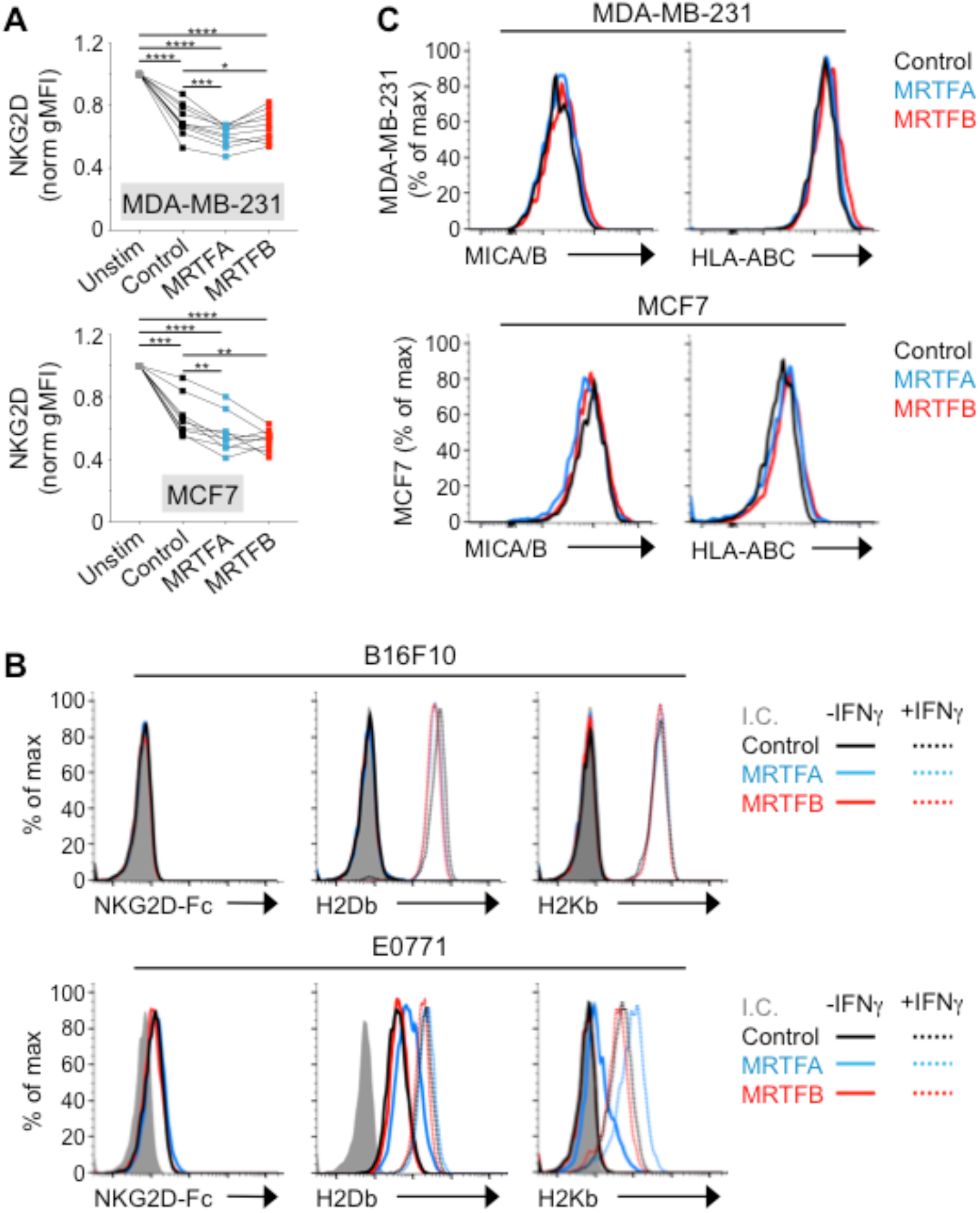
Surface protein expression on cells overexpressing MRTFA or MRTFB. Related to Fig. 6. (A) Flow cytometric analysis of NKG2D expression on human NK cells after 5 h coculture with the indicated target cells. Data were normalized against NKG2D levels on untreated NK cells. Gray lines denote samples derived from the same donor. \*\*\*\**p* < 0.0001, \*\*\**p* < 0.001, \*\**p* < 0.01, and \**p* < 0.05 calculated by one-way ANOVA (*n* = 9 donors for MCF7 experiments, *n* = 10 donors for MDA-MB-231 experiments). Data pooled from 3 independent experiments. (B) Flow cytometric analysis of NKG2D ligands and MHC proteins (H2Db and H2Kb) on the indicated control or MRTFA/B overexpressing B16F10 and E0771 cell lines. Cells were untreated (solid lines) or pretreated overnight with IFN*γ* (dotted lines). Isotype control (I.C.) is shown in gray. Histograms are representative of 3 independent experiments. (C) Flow cytometric analysis of MICA/B (NKG2D ligands) and HLA-ABC (MHC) on the indicated control and MRTFA/B overexpressing MDA-MB-231 and MCF7 cell lines. Histograms are representative of 3 independent experiments.

**Fig. S7.**
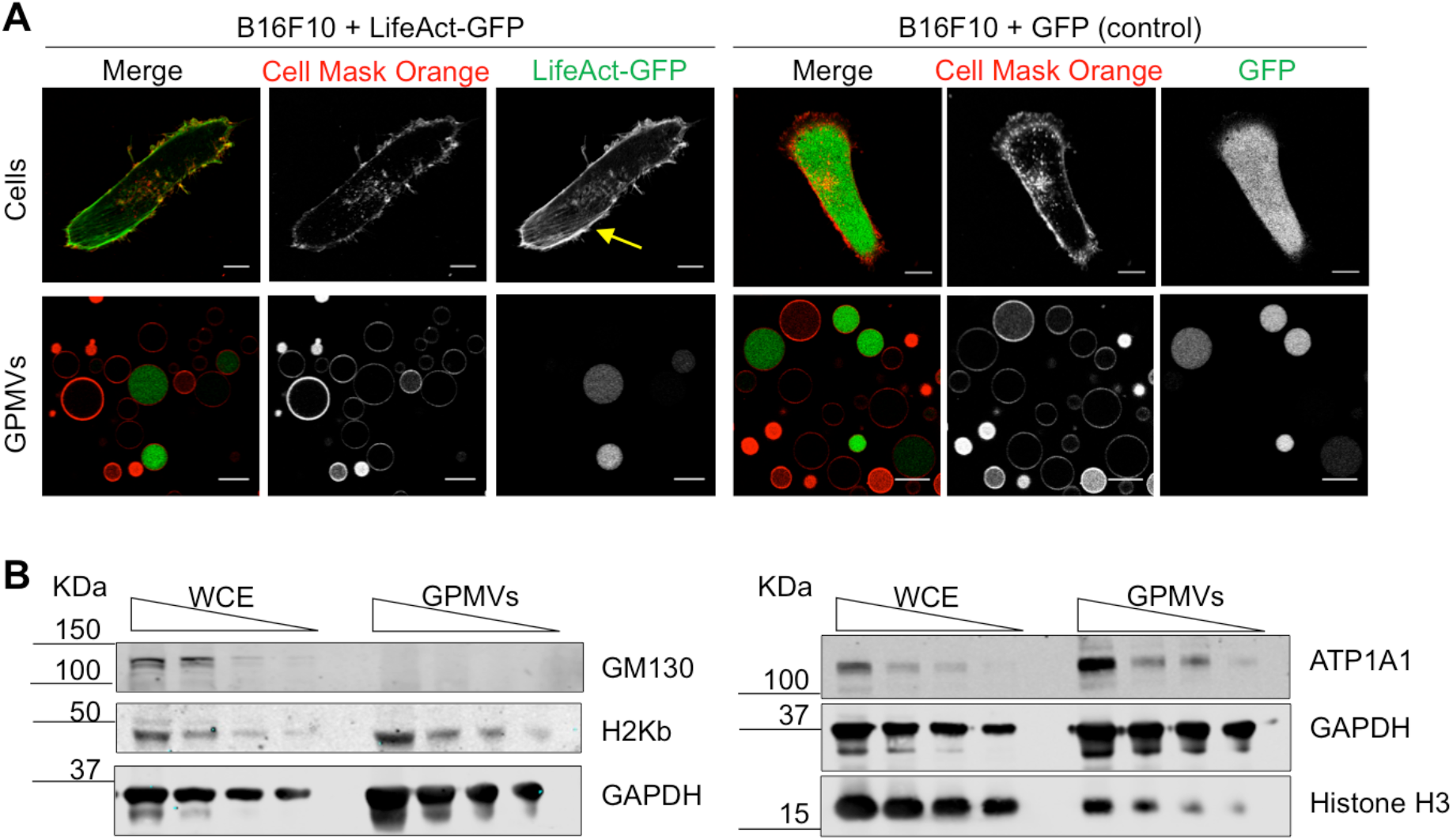
Characterization of giant plasma membrane vesicles (GPMVs). Related to Fig. 7. (A) Representative confocal images of whole cells (above) and cell-derived GPMVs (below). B16F10 cells overexpressing LifeAct-GFP (green, left panel) or GFP control (green, right panel) were labeled with the plasma membrane dye Cell Mask Orange (red). Yellow arrow indicates cortical F-actin present in cells but not in GPMVs. Scale bars: 10 μm. (B) Immunoblots of the indicated proteins, performed using serial dilutions of whole cell extracts (WCE) or GPMVs derived from B16F10 cells. GADPH served as a loading control. Data are representative of 3 independent experiments.

## ACKNOWLEDGEMENTS

We thank C. Firl, L. Stafford, and X. Zheng for technical support; the MSKCC Molecular cytology Core Facility for assistance with imaging and AFM; E. Chan for assistance with image analysis; the MSKCC Integrated Genomics Operation (IGO) and Bioinformatics Core for RNA sequencing and analysis; UIC Research Informatics Core and Z.A. Lei for assistance with TCGA GSEA analysis; K. Pham and A. Rudensky for critical reading of the manuscript; N. Biais, F. Paumet, D. Yuan, L. C. Kam, and members of the M. H., J. M., K. C. H., and J. C. Sun labs for advice and assistance with experiments. Supported by the NIH (R01-AI087644 to M. H., P01-CA94060 to J. M., R01-AI125651 to K. C. H., and P30-CA008748 to MSKCC), the NSF (CMMI-1562905 to M. H.), the Parker Institute for Cancer Immunotherapy (M. H.), NCATS (UL1TR002003 to UIC, RIC), Cycle for Survival (IGO), the Kravis Center for Molecular Oncology (IGO), the Leukemia and Lymphoma Society (M. H.), the Ramón Areces Foundation (M. T.-L.), and UIC (Department of Physiology and Biophysics Funds to E. E. E.).

## AUTHOR CONTRIBUTIONS

M. T.-L., E. E. E., K. S. and M. H. conceived and designed experiments. M. T.-L., E. E. E., K. S., and J. H. collected and analyzed data. Y. R. and A. C. provided critical technical assistance with AFM experiments. M. H., J. M., and K. C. H. supervised research and provided grant support. E. E. E., M. H., M. T.-L., K. S. and J. M. wrote the manuscript.

